# Pyruvate carboxylation identifies Glioblastoma Stem-like Cells opening new metabolic strategy to prevent tumor recurrence

**DOI:** 10.1101/2022.07.18.500427

**Authors:** Ophélie Renoult, Catherine Gratas, Noémie Joalland, Mélanie Laurent--Blond, Hala Awada, Marine Bourgeois, Lisa Oliver, Sophie Chiavassa, Mikaël Croyal, François Paris, Claire Pecqueur

**Author notes:** Corresponding author : Claire Pecqueur CRCI2NA, Nantes Université, INSERM U1307, CNRS 6075, 8 quai Moncousu 44007 Nantes, France. **E-mail**. **Authors contributions** :CP designed research; LO provided primary cultures; OR, CG, NJ, MLB, HA, MB, SC, MC performed research; OR, CG, FP, CP analyzed data; OR, CP wrote the manuscript; all authors revised the manuscript.

## Abstract

Glioblastoma (GBM) are currently associated with a dismal prognosis due to therapeutic resistance. Within the diverse tumor subpopulations, Glioblastoma Stem-like Cells (GSC) have been involved in GBM recurrence. In our study, we demonstrated that these tumor cells can be identified through singular mitochondrial alternative metabolisms. Combining state-of-the-art metabolic studies and the development of a straightforward tumoroid model recapitulating key features of primary GBM cultures, we uncovered a significant use of α-ketoglutarate reductive carboxylation and pyruvate carboxylation in tumoroid GBM cells, catalyzed respectively by isocitrate dehydrogenase and pyruvate carboxylase enzymes. We demonstrated that these singular metabolic features are shared by GBM cells from the mesenchymal subtype and radiation-escaping cells, also involved in recurrence. Finally, we demonstrated that pyruvate carboxylation is required for GBM cell survival in hypoxic niches where glutamine is restricted. Thus, besides providing a new way to identify GSC, our study also opens new therapeutic strategy to limit GBM recurrence.

## Introduction

Glioblastoma (GBM) is the deadliest primary brain tumor in adults. Current standard of care combines surgery, radio- and chemotherapy. However, treatment efficacy remains limited as shown by systematic GBM recurrence and poor patient outcomes (median survival of 18 months and less than 5% of 5-year survival) (Stupp *et al*., 2009). Current clinical obstacles contributing to poor GBM outcome include late diagnosis due to non-specific symptoms, diffuse tumoral infiltration, hidden quiescent states behind the blood-brain barrier, and resistance to conventional therapy. These challenges are further complicated with the intrinsic heterogeneity and the dynamic plasticity of GBM cells, as shown by recent high throughput genomic studies (Patel *et al*., 2014; Meyer *et al*., 2015; Wang *et al*., 2017). Indeed, genetic, epigenetic, and microenvironmental signals greatly influence cellular programs and drive GBM heterogeneity. One main layer of GBM heterogeneity is the developmental state of cells in the tumor. Indeed, GBM contains subsets of glioblastoma stem-like cells (GSCs) that are thought to represent its driving force, possess tumor-propagating potential and exhibit preferential resistance to radiotherapy and chemotherapies (Bao *et al*., 2006; Chen *et al*., 2012). Importantly, identification of GSC is still highly challenging. Indeed, they can reside in various intra-tumoral niches, express diverse markers and evolve with time and treatments (Sottoriva *et al*., 2013; Patel *et al*., 2014; Meyer *et al*., 2015; Dirkse *et al*., 2019). Furthermore, whether GSC markers are expressed in all GSC, identify the same population, or equate to therapeutic response is deeply controversial (Lathia *et al*., 2015; Garnier *et al*., 2019). Functional features are currently used to identify GSC, including a low metabolic state associated with a more quiescent state, the ability to grow as sphere given their self-renewal features, a greater resistance to therapy and the formation of an invasive tumor following orthotopic injection in mice (Chen *et al*., 2010; Tang *et al*., 2021). For decades, cancer research has used diverse established human GBM cell lines growing as adherent monolayers with clonal expansion. These cell lines were easy to manipulate, generated reproducible results and brought significant understanding in GBM biology. However, they did not contain GSC. More recently, development of patient-derived primary cultures has been crucial by providing cell model retaining patient genetic alterations, molecular heterogeneity as well as GSC enrichment when cultured in NSC media (Lee *et al*., 2006). These cell models have tremendously contributed to uncover fundamental mechanisms of cellular GBM drivers depending on patient-specific factors or molecular subtype (Tardito *et al*., 2015; Jin *et al*., 2017; Oizel *et al*., 2017). However, given the great inter- and intra-tumoral GBM heterogeneity, patients-derived primary cultures generate heterogeneous results among cell cultures deriving from different patients, or from different sublocations of the same tumor. While molecular signature identification allow a subsequent distinction between various GSC subpopulations (Jin *et al*., 2017; Oizel *et al*., 2017; Chauvin *et al*., 2019), a significant number of primary cultures is required in order to delineate a global representative and robust biological tumor response. Thus, the development of reliable *in vitro* GBM models, retaining GSC subpopulation and ensuring reliable mechanistic and functional insights, is still required.

Tumoral metabolic adaptations have been demonstrated for decades and belong to current tumor hallmarks (Hanahan and Weinberg, 2011). Importantly, tumor environment shapes tumor metabolic adaptations. Strikingly, relapsing GBM cells, including GSC, are generally located in intra-tumoral hypoxic niches, which are characterized by drastic nutrient unavailability, namely extremely low levels of both oxygen and metabolic substrates (Jin *et al*., 2017). While hypoxic niches limit treatment accessibility and efficacy, they also drive strong tumor metabolic reprogramming. In particular, in addition to canonical glycolysis and oxidative metabolic pathways, hypoxic cells can use carbon skeletons from glycolysis and glutaminolysis through a truncated mitochondrial tricarboxylic acid (TCA) cycle combined with mitochondrial alternative pathways disconnected to mitochondrial oxidative phosphorylation to feed their bioenergetic and biosynthetic needs. Those alternative pathways can involve oxaloacetate production from pyruvate through pyruvate carboxylase (PC) (Cheng *et al*., 2011; Lin *et al*., 2020) or reductive carboxylation of α-ketoglutarate (αKG) into citrate through isocitrate dehydrogenase (IDH) (Mullen *et al*., 2012; Gaude *et al*., 2018). While these alternative metabolic pathways have been investigated in tumor cells with defective mitochondria or under hypoxia, their relevance in GBM biology remains poorly understood.

In the present study, we developed a simple and reproducible GBM cell model at the interface between GBM cell lines and patient-derived primary cells. Using extensive metabolic assays, we uncovered a singular metabolic rewiring in GSC. In particular, we demonstrated that GSC use alternative mitochondrial metabolism to sustain their survival, even in oxygen and nutrient-rich conditions. Interestingly, using diverse *in vitro, in vivo*- and patient-derived cells, we determined that this singular metabolism is shared by all GBM cells involved in GBM recurrence, namely GSC, GBM cells from the mesenchymal subtype and radiation-escaping cells. Finally, we established that genetic and pharmacological inhibition of pyruvate carboxylation significantly decreased GSC survival, in particular following glutamine restriction. Altogether, this work adds valuable knowledge in GBM metabolic flexibility and identify PC as both a new GSC molecular marker and target to reduce GBM recurrence.

## Results

### Implementation of a simple tumoroid model of GBM recapitulating key features of primary GBM cultures

To combine the complexity of GBM biology and the simplicity of an experimental model to ensure reliable mechanistic and functional insights, we investigated the relevance of culturing the human GBM cell line U251 in neural stem cells media (NSC) versus cells cultured in complete medium containing serum (CPT) and to patient-derived primary cultures (cultured in NSC medium). Of note, patient-derived primary cultures were grouped based on their molecular signature assigned based on Verhaak classification (Verhaak *et al*., 2010). As expected, and similarly to primary cultures, after 3 weeks in NSC medium, U251 cells spontaneously grew as tumoroids (**Supplementary 1A**). In contrast, U251 cells in CPT medium grew as an adherent monolayer. Furthermore, the transcriptomic profiles of U251 cells were clearly distinct depending on their culture condition (**Figure 1A**). Indeed, RNA sequencing followed by unsupervised hierarchical clustering discriminated U251 cells cultured in NSC versus CPT medium. As tumoroid structure is associated with GSC enrichment, we performed extreme limiting dilution assay in order to determine the amounts of GSC in the cell cultures (**Figure 1B**). U251 cells cultured in CPT medium did not contain GSC cells. In contrast, U251 cells cultured in NSC medium were enriched in GSC (16.5±2.7%), even if a higher GSC enrichment was observed in primary cultures (38.6 ±14.3%). GSC enrichment was further investigated through expression analysis of several markers associated with either stemness or differentiation. Expression of the stemness marker CD133 was strongly expressed in U251 cells cultured in NSC medium but barely detected in U251 cells in CPT medium (**Figure 1C**). GSC enrichment in NSC condition was further confirmed by the reduced expression of GFAP and β3-TUB, differentiation markers of astrocyte and neuron respectively (**Figure 1C**). Investigation of other stemness markers expression, both at the protein and the RNA levels, did not reveal significant differences between NSC and CPT conditions (**Supplementary 1B**). Importantly, these results were in agreement with the great discrepancy of stemness markers expressions observed between primary cultures (**Supplementary 1C-D**). Expression of markers associated with GBM aggressiveness, usually overexpressed in primary GBM cells from the MES subtype, was also analyzed and revealed that U251 cells cultured in NSC medium displayed higher expression of CHI3L1, ASCL1 and CD109 than their CPT counterparts (**Figure 1C and Supplementary 1C**). To better characterize U251 cells cultured in NSC medium, we further explored how culture conditions affected U251 cell phenotype. U251 cells cultured in NSC medium displayed a reduced proliferation rate compared to those cultured in CPT medium (**Figure 1D**). Their orthotopic injection in mice brain triggered the development of a highly infiltrative tumor as revealed by tumor IHC staining against MHC-I (**Figure 1E, left panel**). Accordingly, a significant delay in clinical signs appearance was observed in mice bearing tumor from U251 cells cultured in NSC medium (**Figure 1F, right panel**). Tumor cell sensitivity to increasing doses of radiation was measured by flow cytometry following PI staining. Interestingly, while increasing doses triggered increasing cell death, U251 cells cultured in NSC medium were more resistant to radiation than those cultured in CPT medium (**Figure 1F**). Finally, we compared the transcriptomic profile of U251 cells with the different primary cultures (**Figure 1G, green vs red**). Interestingly, unsupervised PCA analysis clustered U251 cells cultured in NSC medium with primary GBM cultures, in contrast to U251 cells cultured in CPT medium, which clearly stand outside (**Figure 1G**). Furthermore, U251 cells cultured in NSC medium roughly stood in the middle between the MES and the CPN primary cultures.

**Figure 1:**
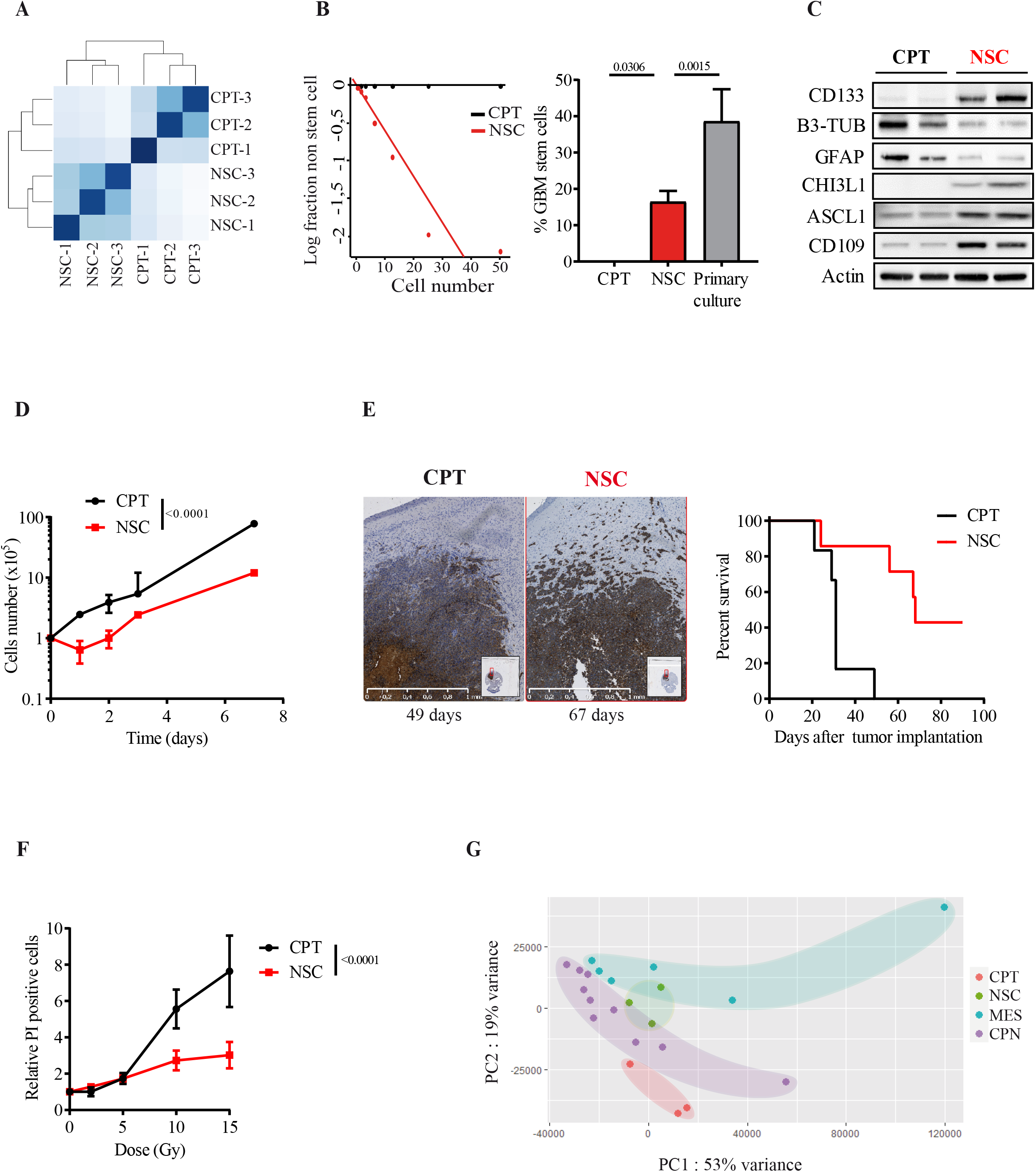
U251 cells cultured in NSC media recapitulate key features of patient-derived primary cultures. **A**. Heatmap of Euclidean distance between U251 cells in CPT versus NSC media. **B**. Stem cell enrichment measured by limiting dilution assay in U251 cells plated in CPT (*black*) and NSC (*red*) media and patient-derived primary cultures. Representative plot of the log fraction of wells without spheres in function of the plated U251 cell number (*left panel*) and subsequent calculated frequency of GSC (*right panel*). One-way ANOVA (n ≥3). **C**. Representative immunoblots of stemness, differentiation and aggressiveness markers expression in U251 cells according to culture conditions. For each culture condition, two independent samples are shown. **D**. Cell proliferation of U251 cells depending on culture conditions over time. Student t-test p<0.0001 (n ≥3). **E**. *In vivo* characterization of orthotopic U251 tumor in immunodeficient mice depending on initial culture condition. Representative picture of orthotopic mice tumor from U251 cells cultured in CPT or NSC media (*left panel*) and mice survival curves (*right panel*). n≥6 mice/group, log-rank test p=0.0462. **F**. Relative cell death measured by flow cytometry following PI staining 72h after irradiation. Two-way ANOVA p<0.0001 (n ≥3). **G**. Unsupervised PCA plot of U251 cells in CPT or NSC media, and patient-derived primary cultures, using the top 5000 differential genes. Depending on their molecular subtypes based on Verhaack classification, mesenchymal and classical/proneural patient-derived primary cells were annotated respectively as MES and CPN.

Altogether, our data demonstrate that U251 cells cultured in NSC medium recapitulate key features of primary GBM cells, namely their transcriptomic profile, growth as 3D-tumoroids, GSC enrichment, *in vivo* infiltration and radiation resistance. From here-after U251 cells cultured in NSC medium will be referred as tumoroids while those cultured in CPT medium will be referred as adherent cells.

### Tumoroid cells are poised into pro-oxidative and low metabolic states

GSC can reprogram their metabolism to flexibly adapt to environmental changes, which is considered critical to sustain their self-renewal capacity and their compensative anti-oxidant state. First, the oxidant state in tumoroid or adherent U251 cells was assessed by cytometry analyses following either CM-H2DCFDA staining or MitoSOX staining. Surprisingly, a significant pro-oxidative state was observed in tumoroids compared adherent cells, as shown by higher levels of both cytosolic and mitochondrial ROS (**Figure 2A-B**). Accordingly, several proteins involved in anti-oxidant responses were overexpressed in tumoroids (**Figure 2C**). Global metabolic analyses were first investigated using the Seahorse technology where oxygen consumption rate (OCR) and extracellular acidification rate (ECAR) were measured. As expected, adherent U251 cells exhibited a metabolic energetic profile characterized by a strong OCR and ECAR. Both OCR and ECAR were significantly reduced in tumoroids, in agreement with their more quiescent state (**Figure 2D**). We further deciphered these glycolytic and mitochondrial metabolic discrepancies through quantification of various metabolites by LC-HRMS (**Figure 2E-J**). First, while no difference in glucose consumption was measured (**Figure 2F**), glutamine consumption was significantly reduced in tumoroid cells as compared to adherent cells (**Figure 2G**). Surprisingly, almost all intracellular metabolites measured were significantly more abundant in tumoroid cells as compared to adherent cells (**Figure 2H-J**). Indeed, key glycolytic metabolites such as pyruvate, lactate and alanine were 2 to 50-fold more abundant in tumoroids than in cells growing as adherent (**Figure 2H)**. Similarly, glutamine, citrate, succinate, fumarate and malate metabolism, which are involved in mitochondrial anaplerosis, were far more abundant in tumoroids than in adherent cells **(Figure 2I-J)**. Of note, glutamate levels were similar in tumoroid and in adherent cells (**Supplementary 2A**).

**Figure 2:**
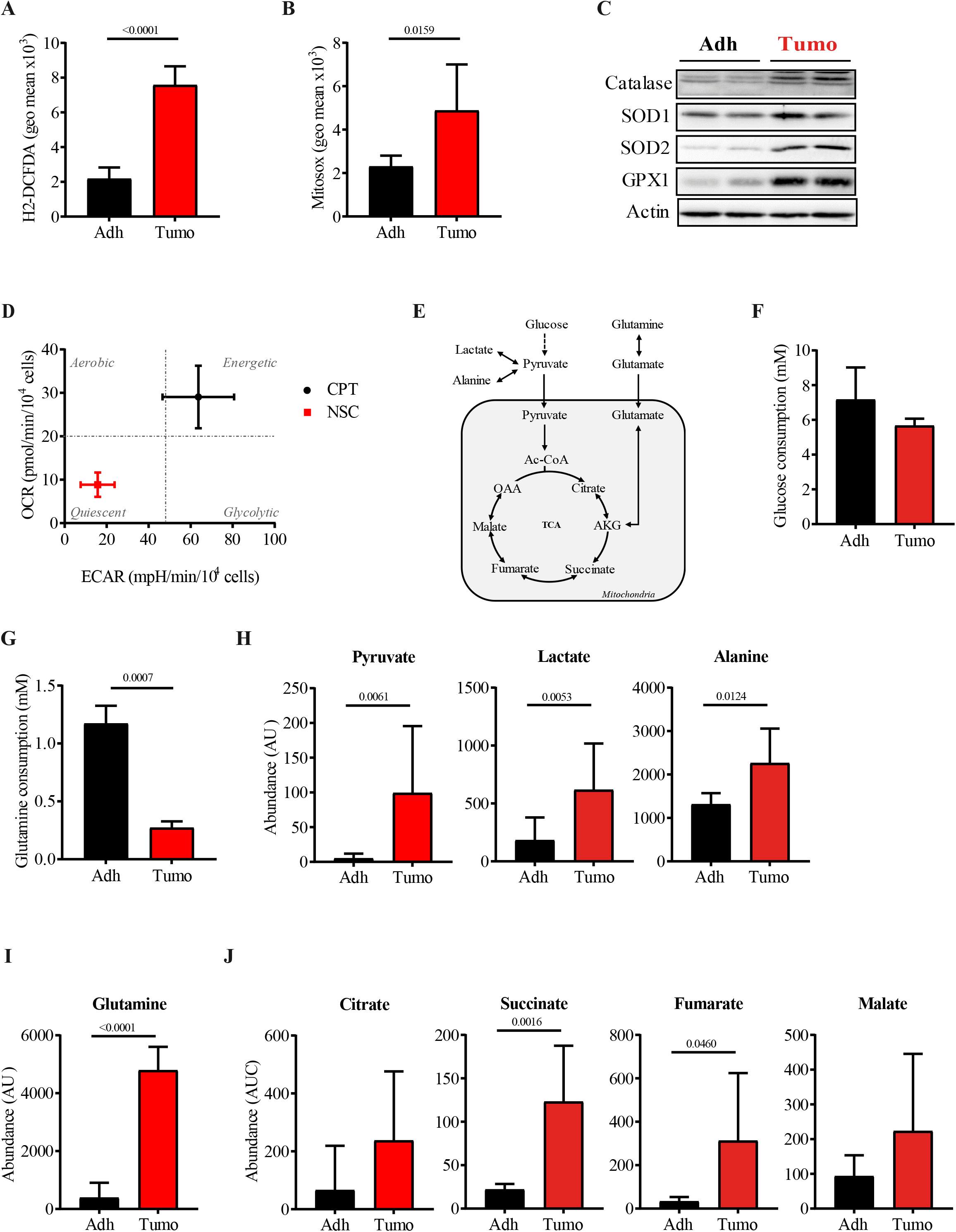
Global metabolic rewiring in tumoroid U251 cells. **A-B**. Total ROS (A) and mitochondrial superoxide (B) levels were measured by flow cytometry after CM-H2DFCDA and Mitosox staining respectively, in adherent (*black*) and tumoroids (*red*) U251 cells. (n=3). **C**. Representative immunoblot of anti-oxidant enzymes in adherent and tumoroid U251 cells. For each tumor population, two independent samples are shown. GPX1: Glutathione Peroxidase-1, SOD1: superoxide dismutase-1, SOD2: superoxide dismutase-2. **D**. Metabolic profile of adherent (*black*) and tumoroids (*red*) U251 cells using the Seahorse technology. Results are presented as Oxygen Consumption Rate (OCR) in function of Extracellular Acidification Rate (ECAR) (n=4). **E**. Schematic glucose and glutamine fueling of mitochondrial metabolism. **F-G**. Glucose (F) and glutamine (G) consumption of adherent (*black*) and tumoroid (*red*) U251 cells over 48 hours. (n=3). **H-J**. Intracellular abundance of key metabolites. Levels of glycolytic metabolites (**H**), glutamine (**I**) and TCA intermediates (**J**) were measured by mass spectrometry over 5 hours. (n=3). Statistical analyses were performed using Student t-test in all experiments.

Thus, our data show that, in contrast to U251 cells growing as adherents, tumoroid cells are poised into a pro-oxidative state associated with a low metabolic state leading to a global accumulation of most key intracellular metabolites.

### Tumoroid cells preferentially use glutamine to sustain their reductive carboxylation rather than the TCA cycle

Our previous data highlight tumor metabolic flexibility. To underpin key metabolic reprograming occurring in U251 cells growing as tumoroids, we further investigated glutaminergic metabolism in these cells. First, supervised analysis of genes involved in mitochondrial anaplerosis using our RNA sequencing data was performed and clearly revealed a distinct transcriptomic pattern between tumoroid and adherent cells (**Figure 3A**). Since glutamine is one of the main mitochondrial anaplerotic substrate, expression of key enzymes involved in glutamine catabolism or synthesis was investigated (**Figure 3B-C**). Adherent U251 cells clearly used glutamine to fuel their mitochondrial metabolism as shown by high level of glutaminase (GLS) to hydrolyze glutamine into glutamate, and undetected glutamine synthase (GS), which catalyzes the opposite reaction. In contrast, tumoroid U251 cells exhibited the exact opposite pattern with high expression of GS and low expression of GLS. Strikingly, the second step of glutamine catabolism toward mitochondrial anaplerosis involve distinct enzymes between cells. While adherent cells expressed more glutamic-oxaloacetic acid transaminase (GOT1), tumoroid cells showed higher level of glutamate dehydrogenase (GDH). We then measured how CB839, a synthetic inhibitor of GLS, would affect OCR in these tumor cells. Surprisingly, CB839 inhibited 39±18.1% of glutamine-driven OCR while it inhibited only 13±3.3% of glutamine-driven OCR in adherent cells (**Figure 3D**). Altogether, these results suggest that tumoroid cells exhibit a complex glutamine metabolism and are using a significant amount of glutamine to fuel their mitochondrial anaplerosis, despite GS overexpression. To confirm this hypothesis, we cultured tumoroid and adherent cells in medium containing ^13^C_5_-glutamine and measured ^13^C enrichment of intracellular metabolites by LC-HRMS. When glutamine fuels mitochondrial anaplerosis, ^13^C_5_-αKG (m+5) is generated from ^13^C_5_-glutamine, subsequently decarboxylated by αKG dehydrogenase (AKGDH) into succinate (m+4) and ultimately leading to the generation of citrate (m+4) through the TCA cycle (**Figure 3E**, blue pathway). AKGDH activity was significantly reduced in tumoroid cells as compared to adherent cells (**Figure 3F**). Instead, they produced higher amount of citrate (m+5) deriving from the direct reductive carboxylation of ^13^C_5_-αKG (m+5) through isocitrate dehydrogenase (IDH) (**Figure 3E-F**, *orange pathway*). In fact, the ratio of αKG reductive carboxylation over TCA cycle was 5-time higher in tumoroid cells than in adherent cells (0.49 ± 0.11 versus 0.11 ± 0.05 respectively)(**Figure 3G**). Finally, to determine whether high ratio of αKG reductive carboxylation over TCA was specific to U251 cells or is widely observed in GSC cultures, we measured these fluxes in MES and CPN primary cultures. Fluxomic analysis following ^13^C_5_-glutamine labeling revealed that the mean ratio of reductive carboxylation over TCA greatly varied between patient-derived primary cultures depending on their molecular subtype, but was globally very similar to the one observed in U251 cells cultured in NSC medium (0.55 ± 0.23)(**Figure 3H**). Indeed, ratio of αKG reductive carboxylation over TCA cycle varies from 0.2 to 0.7 in primary GBM cells from the CPN subtype and from 0.5 to more than 1 in cells from the mesenchymal subtype. In fact, all patient-derived primary cultures exhibited similar enrichment in citrate (m+5) while citrate (m+4) enrichment was significantly lower in MES primary cells than in CPN cells (**Figure 3H**). In contrast to what was observed in U251 cells, MES primary cells with higher ratio of αKG reductive carboxylation over TCA cycle overexpressed both GLS and GDH as compared to CPN primary cells (**Figure 3J**). Furthermore, IDH1 enzyme, which is involved in reductive carboxylation flux, was found overexpressed in CPN primary cells. Thus, the key enzymes expression involved in glutamine metabolism does not correlate with αKG metabolic fate in tumor cells.

**Figure 3:**
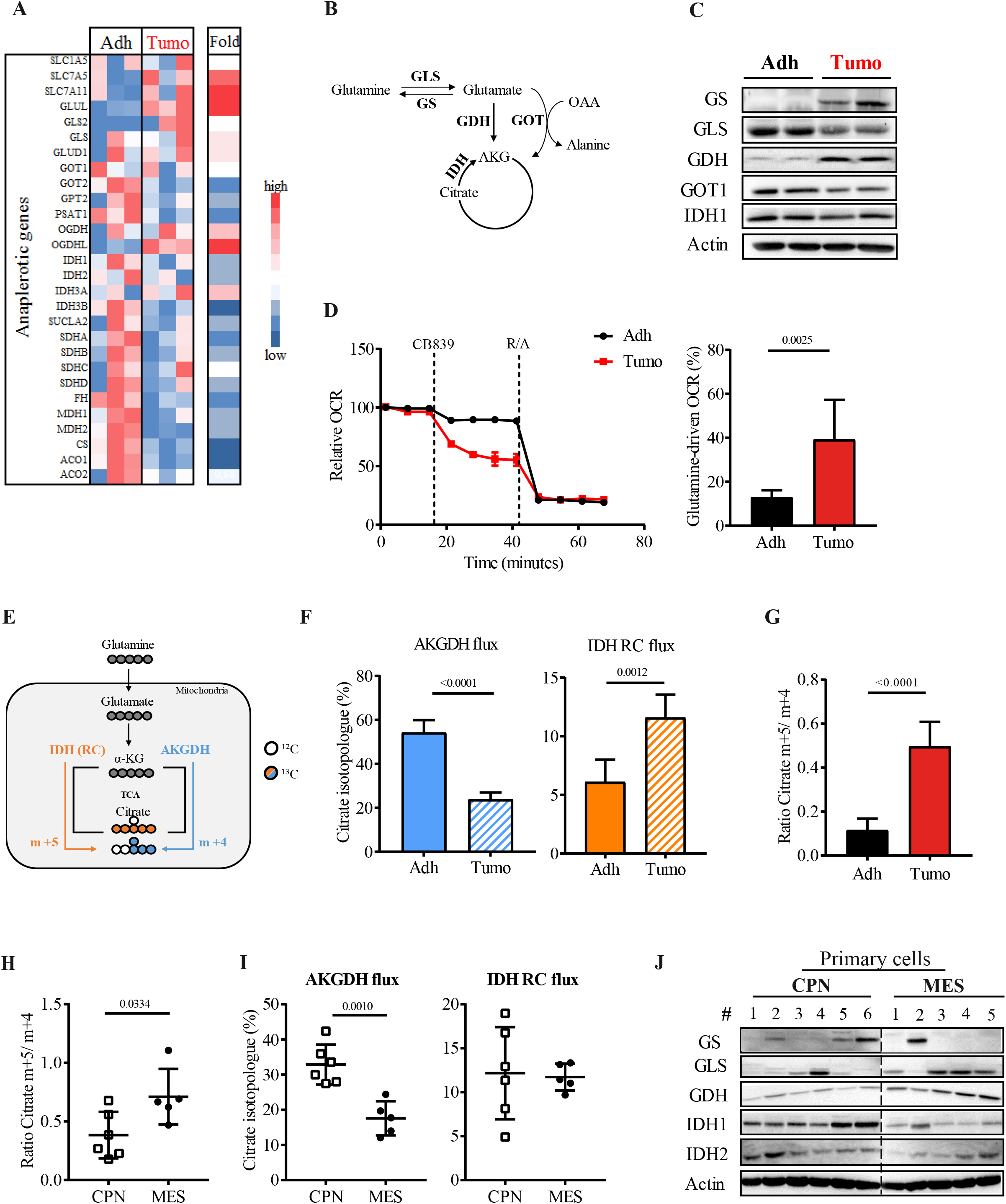
Mitochondrial glutamine metabolism discriminates tumoroid from adherent tumor cells. **A**. Heat map of read number from genes involved in anaplerotic metabolism in adherent and tumoroid U251 cells. Expression fold was calculated as the ratio of gene read number of tumoroid over adherent cells. **B**. Schematic glutamine metabolic pathway sustaining mitochondria metabolism. **C**. Representative immunoblot of main enzymes involved in glutamine metabolism in adherent and tumoroid cells. For each tumor population, two independent samples are shown. GS: Glutamine Synthetase, GLS: Glutaminase, GOT1: Glutamic-oxaloacetic transaminase 1, GDH: Glutamate Dehydrogenase, IDH1: Isocitrate Dehydrogenase-1. **D**. Glutamine-driven oxygen consumption rate (OCR) in adherent and tumoroid U251 cells. Relative OCR in tumor cells monitored with time following injection of GLS inhibitor, CB839 (*left panel*). Percent of glutamine-driven OCR was calculated as the difference between OCR before and after CB839 injection (*right panel*) (n=3 in duplicates). **E**. Schematic mitochondrial glutamine-derived metabolites fate using ^13^C_5_-labeled glutamine. Glutamine-derived (m+5) αketoglutarate (αKG) fuels mitochondrial metabolism through its oxidative decarboxylation catalyzed by AKGDH (*blue)* or its reductive carboxylation (RC) catalyzed by IDH (*orange*), leading respectively to (m+4) or (m+5) citrate enrichment. **F**. AKGDH (*left panel*) and IDH (*right panel*) activities determined by ^13^C-citrate enrichment using mass spectrometry analyses following culture with ^13^C_5_-labeled glutamine. (n=5). **G**. Ratio of ^13^C-citrate isotopologues (m+5) over (m+4) in adherent (*black*) and tumoroid (*red*) U251 cells. Statistical analyses were performed using Student t-test in all experiments. **H**. Ratio of ^13^C-citrate isotopologues (m+5) over (m+4) in CNP (*empty squares)* and mesenchymal (*black circles*). **I**. AKGDH (*left panel*) and IDH (*right panel*) activities in patient-derived primary cultures depending on their molecular subtype. Each dot represents one primary culture (n≥2 in duplicates). **J**. Representative immunoblot of key enzymes involved in glutamine metabolism in primary GBM cultures depending on the molecular subtype. Analysis was performed on 6 CPN and 5 MES independent patient-derived primary cultures. Each lane corresponds to one particular primary GBM culture.

Thus, in contrast to adherent cells, U251 tumoroid cells and patient-derived primary cultures preferentially use glutamine to sustain mitochondrial reductive carboxylation of αKG rather than to fuel the TCA cycle.

### Pyruvate carboxylation is a major metabolic pathway in tumoroid cells

Besides glutamine metabolism, reprogramming of glucose metabolism also occurs in tumoroid cells as shown by increased ECAR (**Figure 2D**) and the distinct transcriptomic pattern of glycolytic genes (**Figure 4A**). Expression of key glycolytic rate-limiting enzymes was comparable in tumoroid and adherent cells (**Figure 4B**). Strikingly, expression of the pyruvate carboxylase (PC), an enzyme also involved in alternative mitochondrial metabolic pathway, was overexpressed in tumoroid cells (**Figure 4B**). We further characterized mitochondrial pyruvate fate by monitoring OCR following addition of UK-5099, an inhibitor of the pyruvate carrier responsible of its import from the cytosol into the mitochondria. UK-5099 reduced the pyruvate-driven OCR by almost 30% in tumoroid cells while it inhibited less than 10% of OCR in adherent cells (**Figure 4C**). To further characterize glycolytic pyruvate fate within the mitochondria, cells were cultured with fully labeled ^13^C_6_-glucose and ^13^C enrichment of intracellular metabolites was measured as previously described (**Figure 4D**). First, no significant difference was observed in glucose-derived pyruvate fate within the cytosol in tumoroid and adherent U251 cells, as shown by similar amount of ^13^C_6_-glucose-derived pyruvate (m+3), lactate (m+3) and alanine (m+3) (**Figure 4E**). Once imported within the mitochondria, glucose-derived pyruvate is usually converted by pyruvate dehydrogenase (PDH) to form acetyl-CoA (m+2), which is subsequently condensed with an unlabeled oxaloacetate (OAA) to generate intermediary metabolites (m+2). PC opens an alternative anaplerotic carbon source to fuel TCA through the direct carboxylation of pyruvate to oxaloacetate (OAA). In this case, ^13^C_6_-glucose-derived pyruvate (m+3) generates OAA (m+3) and subsequent malate (m+3) (**Figure 4D**). Thus, respective PDH and PC activities are given by levels of (m+2) and (m+3) intermediary metabolites. Interestingly, while PDH activity was similar in both tumor populations, PC activity strongly contributed to mitochondrial anaplerosis in tumoroid U251 cells while it was barely detected in adherent cells (**Figure 4F**). To make sure that increased PC activity was not due to glucose access, malate (m+3) enrichment was measured depending on adherent or spheroid conditions, or in presence of various amounts of glucose in U251 cells cultured in NSC medium. Importantly, PC activity was neither impacted by cell culture morphology (adherent versus spheroid), nor by glucose concentration (**Supplementary 3A-B**). Finally, we measured both PDH and PC fluxes in patient-derived primary cells. All primary cultures displayed significant PC activity (**Figure 4G**). However, PC activity was globally higher in MES primary cells than in CPN cells, in agreement with their respective PC expression (**Figure 4H**). No significant difference was observed in other glycolytic enzymes expression.

**Figure 4:**
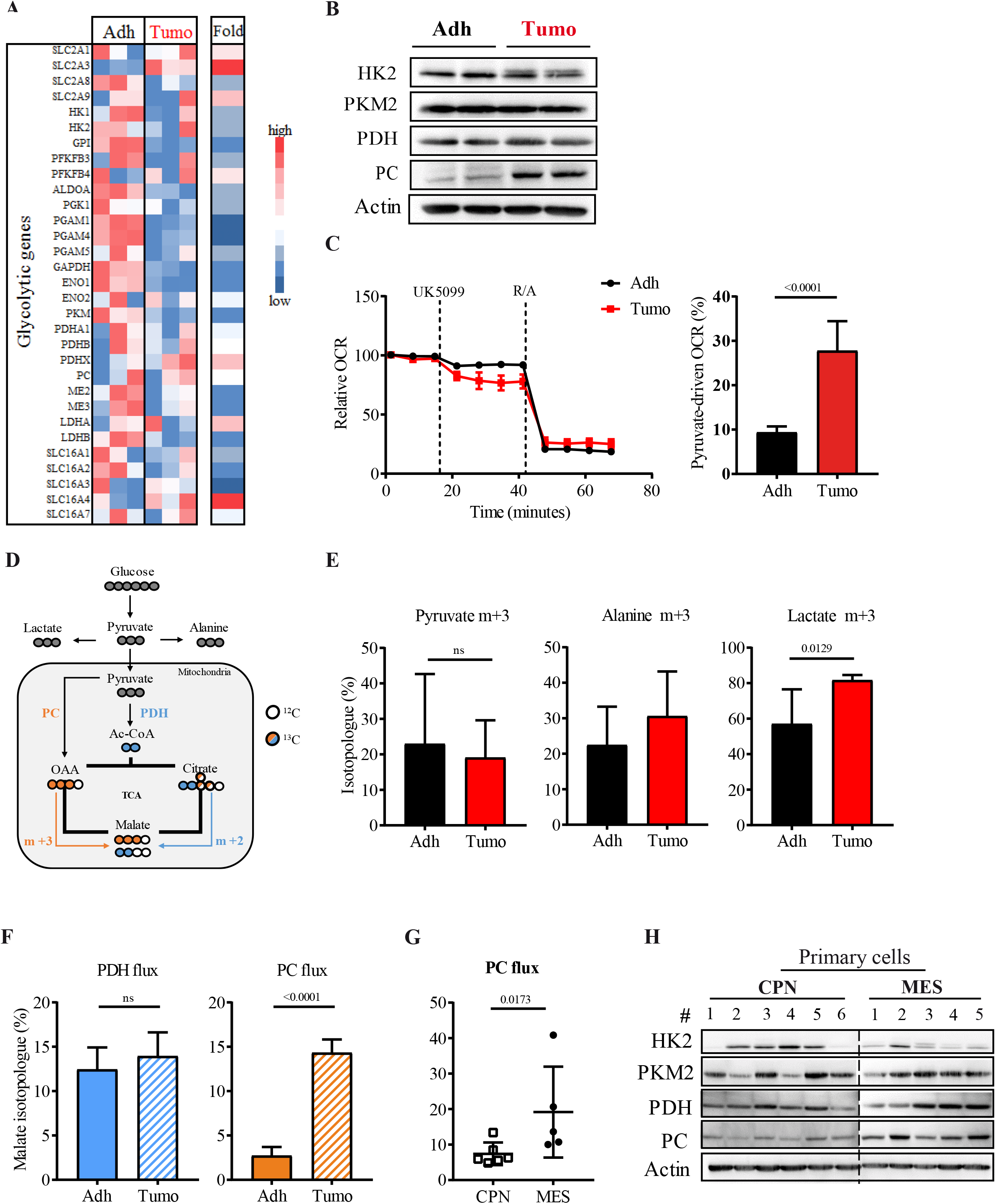
Prominent alternative pyruvate carboxylation in tumoroid U251 cells. **A**. Heat map of read number from glycolytic genes in adherent and tumoroid U251 cells. Fold was expressed as the ratio between tumoroid and adherent gene read number. **B**. Representative immunoblot of key glycolytic enzymes in adherent and tumoroid cells. For each tumor population, two independent samples are shown. HK2: Hexokinase-2, PKM2: Pyruvate kinase M2, PDH: Pyruvate Dehydrogenase, PC: Pyruvate Carboxylase. **C**. Pyruvate-driven oxygen consumption rate (OCR) in adherent and tumoroid U251 cells. Relative OCR in tumor cells monitored with time following injection of mitochondrial pyruvate carrier inhibitor, UK5099 (*left panel*). Percent of pyruvate-driven OCR was calculated as the difference between OCR after and before UK5099 injection (*right panel*) (n=3 in duplicates). **D**. Schematic mitochondrial pyruvate-derived metabolites fate using ^13^C_6_-labeled glucose. Glycolytic pyruvate (m+3) can be either converted to acetyl-CoA (m+2) through PDH or converted to OAA (m+3) by pyruvate carboxylase (PC), subsequently resulting in (m+2) or (m+3) TCA intermediates respectively. **E**. Enrichment of glycolytic metabolites in adherent and tumoroid U251 cells from ^13^C_6_-glucose (n=5). **F**. PDH (*left panel*) and PC (*right panel*) activities determined respectively by (m+2) and (m+3) isotopologues of ^13^C-malate enrichment by mass spectrometry following culture with ^13^C_6_-labeled glucose (n=5). **G**. PC activity in patient-derived primary cultures depending on their molecular subtype. Enzyme activity was determined as ^13^C-malate (m+3) enrichment by mass spectrometry following culture with ^13^C_6_-labeled glucose (n=5). Each dot represents one primary culture (n≥2 in duplicates). **H**. Representative immunoblot of key glycolytic enzymes in primary GBM cultures depending on their molecular subtype. Analysis was performed on 6 CPN and 5 MES independent patient-derived primary GBM cultures. Each lane corresponds to one particular patient-derived primary culture.

Altogether, and similarly to their transcriptomic profiles, U251 cells cultured in NSC medium display a singular use of alternative metabolic pathways, in particular pyruvate carboxylation, which is also observed in patient-derived primary cultures.

### Pyruvate Carboxylase silencing leads to glutamine compensation to sustain mitochondrial metabolism

Since tumoroid U251 cells exhibit significant use of alternative mitochondrial metabolism, we silenced PC and IDH enzymes using the CRISPR-Cas9 technology to investigate their role in tumor cell survival and proliferation. First, silencing of either IDH1 or IDH2, the 2 enzymes able to catalyze reductive carboxylation, specifically decreased citrate (m+5) enrichment by 50% with no effect on citrate (m+4) (**Supplementary 4A-C**). However, IDH1 or IDH2 silencing had no effect on cell proliferation nor cell death (**Supplementary 4D-E**). Similar experiments were then performed following PC silencing using 2 distinct sequence guides (**Figure 5A**). PC silencing was functionally validated using fluxomic analysis following tumor cell culturing with^13^C_6_-glucose. Indeed, enrichments of malate (m+3), fumarate (m+3), aspartate (m+3) and citrate (m+5) were largely decreased in PC-silenced tumoroid cells (**Figure 5B, Supplementary 5A**). Surprisingly, in tumoroids, PC silencing did not affect cell proliferation (**Figure 5C**) but significantly reduced cell viability as shown by increased IP positive cells (**Figure 5D**). As expected, given their low PC expression, PC silencing affected neither cell proliferation, nor cell death of adherent U251 cells (**Supplementary 5B-C**). Next, we determined whether and how tumoroid U251 cells reprogrammed their metabolism to deal with PC silencing. A significant increase in OCR was observed in tumoroid U251 cells which have been silenced for PC (**Figure 5E**). However, neither PDH activity nor the oxidative metabolism of glucose were modified by PC silencing as shown by similar enrichment of (m+2) metabolites following ^13^C_6_-glucose labeling (**Supplementary 5D**). We then investigated whether PC silencing altered glutamine-derived mitochondrial anaplerosis. While there was no difference in mitochondrial alternative flux through IDH, a significant increase in AKGDH activity was observed in tumoroid U251 cells (**Figure 5F**). Finally, we treated U251, MES or CPN tumoroids with 3-MCPD, a pyruvate analogue which cannot be further metabolized by PC. As revealed by the bright-field pictures, 3-MCPD clearly reduced cell density of U251 tumoroid without affecting their diameter (**Figure 5G, top panel**). Similar experiments performed in presence of the DEVD-GFP probe to monitor caspases cascade activation revealed that the apoptotic pathway was activated following 3-MCPD treatment (**Figure 5G, bottom panel**). The effect of this compound was also addressed in tumoroids obtained with fluorescent patient-derived primary cultures (**Figure 5H and Supplementary 5E**). In these models, fluorescence intensity was measured using the IncuCyte technology to monitor cell survival and proliferation in tumoroids. Similarly to U251 tumoroids, 3-MCPD significantly reduced cell survival and/or proliferation in MES and CPN tumoroids, as shown by the reduced fluorescence (**Figure 5H and Supplementary 5E**). As expected based on their respective PC expression and activity, its effect was stronger on MES tumoroid than in CPN tumoroids. Altogether, our data revealed that PC inhibition disrupted mitochondrial metabolic homeostasis, increased glutamine-derived anaplerosis in tumoroids (**Figure 5K**), and more importantly reduced tumor cell progression.

**Figure 5:**
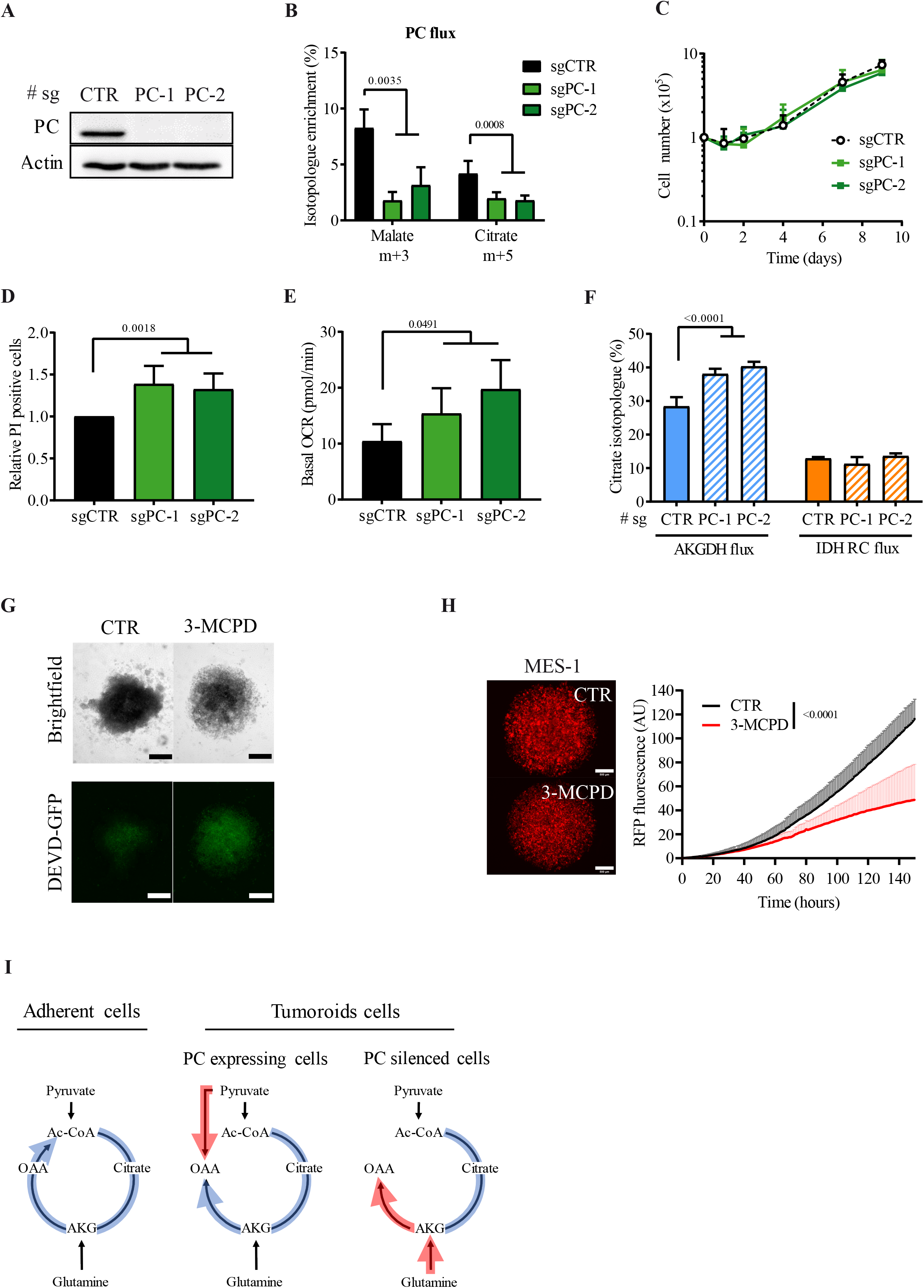
PC inhibition decreases GSC survival. **A**. Representative immunoblot of PC silencing in tumoroid U251 cells. PC was silenced with the CRISPR/Cas9 technology using 2 distinct strand guides (sg). **B**. PC activity determined by isotopologue enrichment of ^13^C-malate (m+3) and ^13^C-citrate (m+5) by mass spectrometry following culture with ^13^C_6_-labeled glucose (n=4). **C**. Cell proliferation of tumoroid cells following PC silencing. Cell proliferation was assessed over 9 days by manual cell counting following eosin staining (n=3). **D**. Relative cell death measured by flow cytometry following PI staining after PC silencing. Two-way ANOVA (n=3). **E**. Basal oxygen consumption rate (OCR) of tumoroid U251 cells following PC silencing (n=3). **F**. AKGDH and IDH RC activities were determined by (m+4) and (m+5) citrate enrichment respectively, from ^13^C_5_-labeled glutamine, using mass spectrometry analyses (n=4). **G**. Cell death after PC pharmacological inhibition with 3-MCPD (7,5mM) in tumoroid U251 cells. DEVD-GFP probe allows apoptosis detection through monitoring of caspases cascade activation leading to GFP emission (Objective 10X, scale bar = 500µm). **H**. Proliferation of RFP-MES-1 primary GBM cells following 3-MCPD treatment (7,5mM). Fluorescence intensity was measured over 8 days with an IncuCyte device (Objective 4X, scale bar = 500µm). Two-way ANOVA (n=3). **I**. Schematic representation of mitochondrial fluxes rewiring in adherent versus tumoroid cells, according to PC expression. Statistical analyses were performed using Student t-test in all experiments, except indicated otherwise.

### Mitochondrial alternative fluxes are a metabolic characteristic of GBM cells involved in recurrence

One main feature contributing to poor GBM outcome is intrinsic therapeutic resistance. We thus investigated whether mitochondrial alternative fluxes were also used in GBM cells escaping to treatment, in particular radiation therapy. Two patient-derived primary cultures from each subtype were submitted to 4 cycles of 5 Gy-radiation every 10 days (20 Gy total) (**Figure 6A**). First, we were able to amplify radiation-escaping cells from both MES primary GBM cultures whereas only one CNP culture over the 2 tested was able to escape radiation and proliferate. Interestingly, fluxomic analysis revealed that radiation-escaping cells globally exhibited an increase in both IDH and PC activities (**Figure 6B-C**). However, while increased in αKG reductive carboxylation occurred only in mesenchymal tumor cells, increased pyruvate carboxylation was observed in all radiation-escaping cells. Of note, neither AKGDH nor PDH activities were altered in radiation-escaping cells (**Supplementary 6A-B)**. Finally, radiation-escaping GBM cells were generated from an orthotopic mouse model bearing mesenchymal GBM and submitted to one single dose of stereotactic radiation (15Gy) (**Figure 6D**). Similarly, to our *in vitro* results, a significant increase of IDH and PC activity was observed in these *ex-vivo* radiation-escaping cells (**Figure 6E-F)**. Again, no difference was observed in AKGDH or PDH activities (**Supplementary 6C-D)**

**Figure 6:**
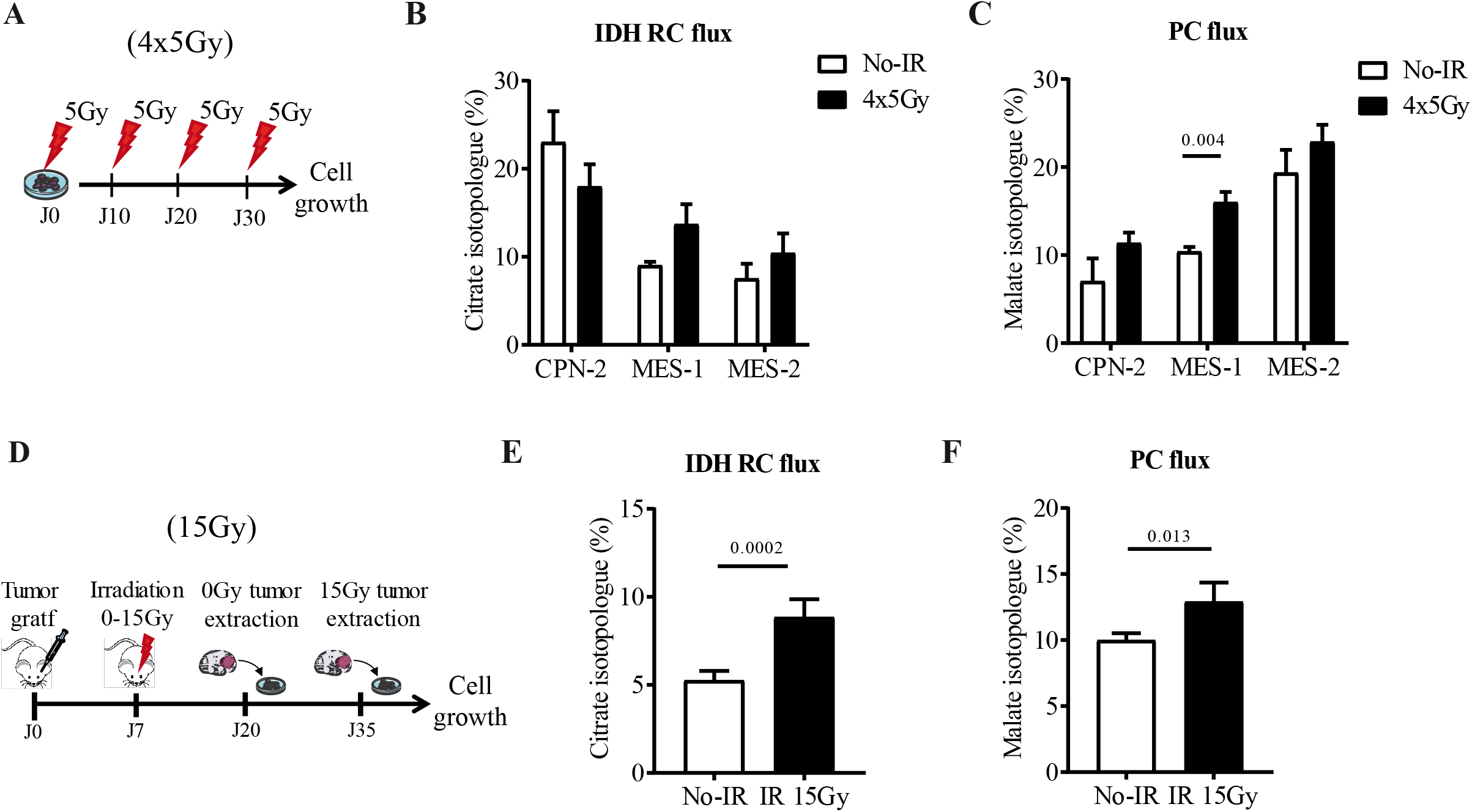
Radiation-escaping cells display increased alternative metabolic fluxes. **A**. Experimental design of *in vitro* generation of radiation-escaping cell following 4 cycles of 5Gy-radiation (4×5Gy). Experiments were performed on 2 CPN and 2 MES independent patient-derived primary cultures. **B-C**. IDH RC (B) and PC (C) activity determined respectively by (m+5) citrate and (m+3) malate enrichment in 4×5Gy radiation-escaping cells (n=3). **D**. Experimental design of *in vivo* generation of 15Gy radiation-escaping cells following stereotactic 15Gy-radiation of the brain hemisphere bearing the tumor in an orthotopic GBM model. 15Gy radiation-escaping cells were isolated from 3 independent brain tumors using MES-1 cells. **E-F**. IDH RC (E) and PC (F) activities were determined respectively by (m+5) citrate and (m+3) malate enrichment in *ex-vivo* 15Gy radiation-escaping cells (n=3 in duplicate).

Altogether, our results revealed that all tumor cells involved in GBM recurrence, namely GSC, mesenchymal cells and radiation-escaping cells, can be identified through their increased alternative mitochondrial fluxes, in particular pyruvate carboxylation.

### PC expression is required when glutamine availability is restricted

GSC and/or mesenchymal GBM cells have been described to be located in intra-tumoral hypoxic niches, which are characterized by lower oxygen and substrate availabilities. First, we investigated by limiting dilution assay whereas PC silencing affected GSC enrichment using tumoroid U251 cells. Interestingly, both sequence guides led to a decrease in GSC numbers (**Figure 7A**). These results were confirmed by immunoblotting since expression of several differentiation markers were increased in these cells as compared to the CTR condition (**Figure 7B**). Then, tumor cell survival was monitored with time following either hypoxia (1% oxygen) or glutamine depletion. Hypoxia reduced cell proliferation in all tumoroid U251 cells, independently of PC expression (**Figure 7C**). In contrast, while glutamine deprivation significantly reduced cell proliferation of PC-expressing cells, U251 cells silenced for PC were not able to survive in absence of glutamine (**Figure 7D**). Metabolic glutamine dependency of PC-silenced cells was further confirmed by Seahorse analysis showing a significant decrease of OCR in absence of glutamine as well as an increased glutamine-driven OCR in its presence (**Figure 7E-F**). Since *in vivo* hypoxic niches probably contained some glutamine, we investigated whereas not only glutamine depletion but also glutamine restriction might alter tumoroid U251 survival. Interestingly, PC-silenced cells exhibited increasing cell death proportionally to increasing glutamine restriction (**Figure 7G**). Finally, to investigate whereas a pharmacological targeting of glutamine metabolism can kill tumoroid U251 cells silenced for PC, cells were cultured in presence of CB839, an inhibitor of glutaminase involved in the first step of glutamine catabolism. While viability of PC-expressing cells viability was not altered by CB-839, PC-silenced cells were not able to survive even at low doses (0.5µM) of CB839 (**Figure 7H**).

**Figure 7:**
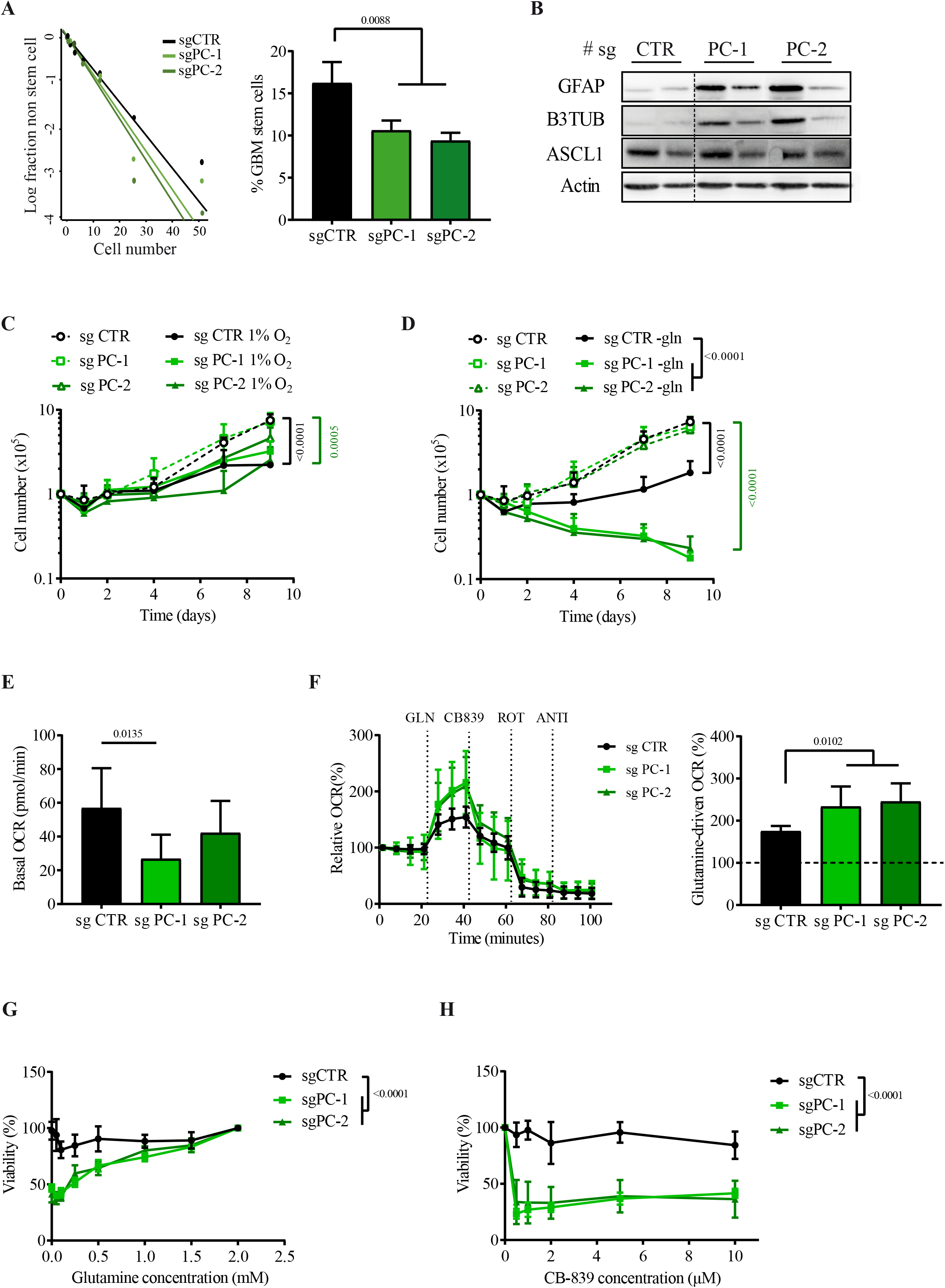
PC is required for GSC survival and following glutamine-restriction. **A**. Stem cell enrichment measured in control (*black*) and PC-silenced (*green*) cells by limiting dilution assay. Representative plot of the log fraction of wells without sphere in function of the plated cell number (*left panel*) and subsequent frequency of GSC (*right panel*). One-way ANOVA (n=3). **B**. Representative immunoblots of differentiation marker expression in tumoroid U251 cells according to PC expression. For each condition, two independent samples are shown. **C**. Cell proliferation of tumoroid U251 cells according to PC expression and oxygen concentration. Cell proliferation was assessed over 9 days in normoxia (*dashed lines*) or hypoxia (1% O_2_, *solid lines*) by manual cell counting following eosin staining. Two-way ANOVA (n=4). **D**. Cell proliferation of tumoroid U251 cells according to PC expression and glutamine concentration. Cell proliferation was assessed over 9 days in complete media (*dashed lines*) or without glutamine (*solid lines*) by manual cell counting following eosin staining. Two-way ANOVA p<0.0001 (n=4). **E**. Basal OCR in glutamine deprived cells according to PC expression. One-way ANOVA (n=3). **F**. Glutamine-driven oxygen consumption rate (OCR) depending on PC expression. Relative OCR monitoring in CTR (*black*) and PC-silenced (*green*) cells when cultured in absence of glutamine for 1 hour. OCR was monitored following successive injections of glutamine (2mM), CB839 (20µM), Rotenone (1µM) and Antimycin A (1µM) (*left panel*). Percent of glutamine-driven OCR was calculated as the difference between OCR before and after glutamine injection (*right panel*). One-way ANOVA (n=4). **G**. Relative cell viability according to PC expression and glutamine concentration. Viability was assessed by MTT staining after 96h of treatment. Student t-test (n=3). **H**. Dose response of GLS inhibitor CB839 in CTR and PC-silenced cells. Viability was assessed by alamar blue staining after 96h of treatment. Student t-test (n=3).

Therefore, pyruvate carboxylation is required for tumoroid U251 cells survival in intra-tumoral niches where glutamine is restricted.

## Discussion

GBM are currently associated to a very poor prognosis due to systematic tumor escape to treatment. Within the various phenotypic and molecular tumor subpopulations, GSC, mesenchymal and radiation-escaping cells, have been involved in GBM recurrence. In our study, we uncovered that all these tumor subpopulations can be identified through their singular metabolic utilization of mitochondrial alternative pathways. Indeed, with the development of a simple, straightforward and reproducible tumoroid model at the interface between human GBM cell lines and patient-derived primary cultures, we demonstrated a significant use of αKG reductive carboxylation and pyruvate carboxylation catalyzed respectively by IDH and PC enzymes these tumor cells involved in GBM recurrence.

One of the main current challenges to study GBM is to find an optimal cell model that offers a fine equilibrium between the complexity of GBM biology and the simplicity of an experimental model to uncover reliable mechanistic and functional insights. Classical *in vitro* GBM models mostly involved established GBM cell lines and patient-derived primary cultures. While established cell lines are easy to manipulate and generate reproducible results, one main limitation is the lack of GSC since serum causes their irreversible differentiation. In contrast, patient-derived primary cells retain GSC subpopulation but are also highly heterogeneous since they are deriving from different tumor sublocations and/or different patients. The main difference in the *in vitro* propagation of these 2 cellular models is serum requirement. Indeed, all established GBM cell lines are grown in media containing serum, whereas patient-derived primary cultures are grown in serum-free media. In our study, we developed a simple and reproducible tumoroid model at the interface between human GBM cell lines and patient-derived primary cultures. Importantly, our model recapitulates key features of patient-derived primary cells, namely growth as tumoroids, quiescence, GSC enrichment, *in vivo* infiltration, radiation resistance as well as metabolic flexibility. Thus, by providing rapid, robust and reproducible results, our tumoroid model appears as an interesting complementary cell model that can be used upstream of the precious and limited patient-derived primary cultures.

In the past decade, several publications have reported alternative mitochondrial flux through αKG reductive carboxylation in tumor cells as a metabolic adaptation of mitochondrial impairment (Metallo *et al*., 2011; Mullen *et al*., 2012; Gaude *et al*., 2018). Reductive carboxylation is the reverse reaction of the conventional clockwise step of oxidative decarboxylation and is exclusively catalyzed through IDH1 and IDH2 enzymes (Alzial *et al*., 2022). Several recent publications revealed the importance of this alternative metabolic pathway for *de novo* lipogenesis in cells exhibiting mitochondrial dysfunction or upon hypoxia (Metallo *et al*., 2011; Wise *et al*., 2011; Gaude *et al*., 2018). In contrast, to our knowledge, only few publications have reported pyruvate carboxylation as a tumor metabolic adaptation (Fan *et al*., 2009; Cheng *et al*., 2011; Phannasil *et al*., 2015; Sellers *et al*., 2015; Davidson *et al*., 2016; Ma *et al*., 2016; Lao-On *et al*., 2020; Lin *et al*., 2020). Furthermore, most of these publications were limited to non-small cell lung (NSCL) and breast cancers and correlated PC expression to tumor aggressivity or metastasis. In our study, we demonstrated for the first time that both alternative mitochondrial pathways were significantly used in GSC, despite functional mitochondria and full availability of both nutrients and oxygen. In agreement with previous studies, expression and function of PC in GBM subpopulations are correlated with tumor cell aggressiveness and ability to escape treatment (Fan *et al*., 2009; Cheng *et al*., 2011; Phannasil *et al*., 2015; Sellers *et al*., 2015; Davidson *et al*., 2016; Ma *et al*., 2016; Lao-On *et al*., 2020; Lin *et al*., 2020). Since GSC reside in hypoxic niches within the tumor (Garnier *et al*., 2019), our results raised the hypothesis of a metabolic imprinting of GBM cells by hypoxia. Indeed, upon hypoxia, combined utilization of αKG reductive carboxylation and pyruvate carboxylation favors the running of TCA cycle in a truncated manner allowing biosynthesis of fatty acid, fueling of anti-oxidant pathways while saving energy.

Therapeutic improvement in GBM treatment clearly relies on the elimination of GSC, mesenchymal GBM cell or/and radiation-escaping cells. While these 3 subpopulations are not mutually exclusive, their clinical identification is currently challenging given the lack of universally informative biomarkers and GBM heterogeneity dynamics. For example, current biomarkers used to identify GSC includes CD133, ALDH, CD90, Nestin or CD44 (Singh *et al*., 2004; Bao *et al*., 2006). However, several studies have failed to confirm their accuracy and reported that mesenchymal GSC cells do not express CD133 while some CD133+ cells are differentiated (Wang *et al*., 2008; Chen *et al*., 2010; Lottaz *et al*., 2010). In our study, we demonstrate that GSC, mesenchymal GBM cells and radiation-escaping cells significantly overexpressed PC. Our results are of significant interest since with the development of high-throughput transcriptomic profiling at the single cell level, recent studies uncovered that GSC and/or molecular subtype are not clonal entities defined by specific functional properties and transcriptomic profiles, but rather correspond to plastic states that most tumor cells can adopt depending on their specific microenvironment (Dirkse et al. 2019; Schmitt et al. 2021; Garofano et al. 2021). In this context, our study provides both a new way to both identify these tumor subpopulations involved in GBM recurrence as well as a new therapeutic strategy to kill them. Indeed, in healthy tissues, PC expression is mostly restricted to liver and β-pancreatic cells (Cappel *et al*., 2019). Furthermore, a recent study reveals that its pharmacological inhibition reduced breast tumor progression both *in vitro* and *in vivo* (Lin *et al*., 2020). Since PC silencing results in increased glutamine dependency, a potential therapeutic strategy against GBM might involve a combination of PC and GLS inhibition in order to obtain significant results.

In conclusion, our study uncovered a metabolic identification of GSC, mesenchymal and treatment-escaping GBM cells that can be exploited to open new targeting strategy to prevent GBM recurrence.

## Material and methods

### Human GBM cells

Human GBM U251MG cell line from ATCC were cultured either in complete media (CPT) composed of DMEM high glucose (21969, Gibco) supplemented with 10% FBS, or in Neural stem cell media (NSC) based on DMEM-F12 supplemented with 50X B-27 (17504-044, Gibco), 100X N-2 (17502-048, Gibco), 25ng/mL bFGF, 20ng/mL EGF, 2µg/mL heparin. Both media were also supplemented with 2mM glutamine (25030-024, Gibco), 100 U/mL penicillin and 100 mg/mL streptomycin (15140-122, Gibco). Primary GBM culture, derived after mechanical dissociation from high-grade glioma as previously described (Oizel *et al*., 2017), were cultured in NSC media. All procedures involving human participants were in accordance with the ethical standards of the ethic national research committee and with the 1964 Helsinki declaration and its later amendments or comparable ethical standards. Informed consent was obtained from all individual participants included in this study. MES-1 and CPN-2 primary cultures were used as fluorescent spheroid. Briefly, 6.10^4^ were plated in poly-HEMA coated 96wells plates (12mg/mL Sigma-Aldrich). Plates were centrifuged at 270g for 3min in order to form spheroids. All cells were maintained at 37°C in 5% CO_2_ atmosphere and regularly checked for mycoplasma contamination. When indicated, cells were irradiated from 0 to 15Gy 24h after plating, at a rate of 1.6Gy/min using Faxitron CP160 (Faxitron X-ray Corporation).

### CRISPR-Cas9 and fluorescent engineering GBM cells

Single-guide RNA sequences were designed using CRISPR design tool (http://crispor.tefor.net) and cloned in the lentiCRISPRv2 plasmid (kindly provided by L. Maillet) (Table 1). LCV2-GFP and LCV2-RFP plasmids were provided by N. Bidère. LentiCRIPSRV2, LCV2-GFP or LCV2-RFP were mixed with sPAX2 and MD2.G plasmids and transfected in HEK293 cells using Lipofectamine 2000 (Life Technologies). After 24h, HEK293 supernatants were collected and added to tumor cell for 48h. Transduced tumor cells were then selected with puromycin (2µg/mL). Protein silencing was confirmed by immunoblot.

**Table 1:**
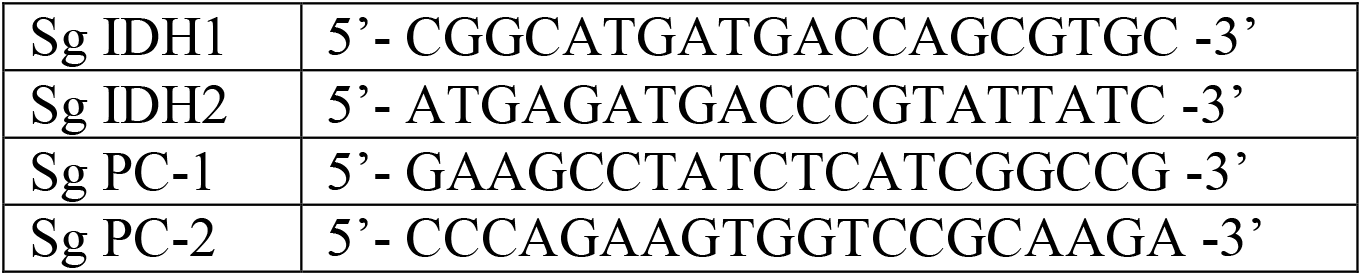
Single guide RNA sequences targeting IDH and PC genes

### 3’ Sequencing RNA Profiling

Total RNA was obtained using the RNeasy mini kit (Qiagen) according to the manufacturer’s instructions. RNA quality was verified with the Bioanalyzer System (Agilent Technologies) using the RNA Nano Chip and analyzed by 3’ Sequencing RNA Profiling (3’SRP). Digital gene expression profiles were generated by counting the number of UMIs associated with each RefSeq genes, for each sample. R package DESeq2 (Bioconductor) was used to normalize gene expression, and detect differentially expressed genes (raw and normalized data deposit on GEO database is in progress). Gene Set Enrichment Analysis (GSEA) was performed using R studio 4.1.1 packages.

### Cell proliferation and viability test

Cells were seeded in NSC media with 0 or 2mM glutamine, in 6 well plate. When indicated, cells were incubated in normoxic or hypoxic conditions (20% and 1% O_2_ respectively) and 1mL of fresh media were added on day 4. For each indicated end-point, cells were mechanically dissociated and counted following eosine-staining.

For dose-response experiment, cellular viability was assessed using MTT or Alamar blue assay as described by the manufacturer. After 2h staining incubation at 37°C, cells were lysed by adding 10% SDS for 4h. Absorbance at 570 and 620nm was measured to calculate cell viability. Spheroids viability of MES-1 and CPN-2 cells was assessed by RFP and GFP quantification respectively. Briefly, spheroids were treated with 7.5mM 3-MCPD over 8 days (Sigma-Aldrich). RFP (625-705nm) and GFP (544nm) fluorescences were acquired with Incucyte system (Sartorius) each hour over 8 days.

### Immunoblot

Protein expression was assessed by immunoblot analysis. Briefly, proteins were obtained following cell lysis with buffer composed of 50mM NaCl, 0.25% DOC, 1% NP40, 0.1% SDS, 1mM EGTA, 1X protease inhibitor cocktail, 50mM Tris HCl pH 7.5. Following an initial saturation step, PVDF membranes were incubated with the primary antibody overnight at 4°C according to the manufacturer recommendations (Table 2). HRP-coupled secondary antibodies (115-035-006 and111-035-006 Jackson ImmunoResearch laboratories) were incubated 1h at room temperature. ECL (WBKLS0500, Millipore) was used for revelation using Fusion-FX Spectra imager (Vilber). While at least three independent experiments were performed, one representative membrane is presented.

**Table 2:** Primary antibody references

| Proteins | Size (kDa) | Supplier | Reference | Dilution |
| --- | --- | --- | --- | --- |
| Actin | 42 | Millipore | MAB1501 | 1/10000 |
| ASCL1 | 78 | Cell Signaling | 4047 | 1/1000 |
| Catalase | 60 | Abcam | 16731 | 1/2000 |
| CD109 | 170 | Santa Cruz Biotechnology | 271085 | 1/1000 |
| CD133 | 133 | Cell Signaling | 64326 | 1/1000 |
| CD44 | 80 | Cell Signaling | 3570 | 1/1000 |
| CH3L1 | 42 | Abcam | 77528 | 1/1000 |
| GDH | 60 | Abcam | 168352 | 1/1000 |
| GFAP | 50 | Cell Signaling | 3670 | 1/1000 |
| GOT1 | 46 | Santa Cruz Biotechnology | 376140 | 1/1000 |
| GLS | 60 | Abcam | 156876 | 1/1000 |
| GPX1 | 22 | Novus Biologicals | NBP1-95368 | 1/2000 |
| GS | 42 | Abcam | 16802 | 1/1000 |
| HK2 | 105 | Cell Signaling | 2867 | 1/1000 |
| IDH1 | 47 | Atlas Antibody | HPA 035248 | 1/1000 |
| IDH2 | 47 | Novus Biologicals | NBP2-43572 | 1/1000 |
| Nestin | 207 | Cell Signaling | 33475 | 1/1000 |
| OLIG2 | 40 | Atlas Antibody | HPA 003254 | 1/1000 |
| PC | 135 | Atlas Antibody | HPA 043922 | 1/1000 |
| PDH | 43 | Cell Signaling | 3205 | 1/1000 |
| PKM2 | 60 | Cell Signaling | C103A3 | 1/1000 |
| SOD1 | 18 | Cell Signaling | 4266 | 1/1000 |
| SOD2 | 25 | Abcam | 13533 | 1/1000 |
| SOX2 | 35 | Cell Signaling | 3579 | 1/1000 |
| β3-TUB | 55 | Cell Signaling | 4466 | 1/1000 |

### Flow cytometry

Total ROS and mitochondrial superoxide were evaluated using the CM-H2DCFDA and mitoSOX red probes respectively (Life Technologies). Briefly, 1×10^5^ cells were seeded in 96 well plates, washed with room temperature PBS, and incubated with the fluorescent probe (2.5µM) for 20min at 37°C. Cell death was investigated following incubation with propidium iodide (PI) (0,4µg/µL) in cold PBS for 15min at 4°C. Fluorescence was acquired on Accuri C6 cytometer (BD Biosciences) and data analysis was performed with FlowJo V10 software.

### Extreme Limit Dilution Assay (ELDA)

To evaluated GSC enrichment, sphere formation was investigated by extreme limiting dilution assay (ELDA) at cell number from 0.3 to 100 cells per well (http://bioinf.wehi.edu.au/software/elda/). Wells were scored as positive if they contained at least one sphere 2 weeks after plating.

### Seahorse analysis

Cells were seeded in XFp plates (Agilent) at either 1.5×10^3^ cells/well the day before experiment when growing in CPT medium, or at 3×10^4^ cells/well on cell tak-coated well (354240, Corning) 1h before experiment when growing in NSC medium. All cells were equilibrated for 1h at 37 °C in bicarbonate-free DMEM supplemented with indicated metabolic substrates. Basal Oxygen Consumption Rate (OCR) and Extracellular Acidification Rate (ECAR) were measured using the XFp Analyzer (Agilent). Glutamine or pyruvate-driven mitochondrial respiration was calculated as the difference between OCR after CB-839 (40µM) or UK5099 (2µM) injection and OCR after rotenone and antimycine A injection (R/A; 1µM each). To evaluate mitochondrial bioenergetics, OCR was measured following the mitostress kit sequence.Data analyses were performed using Seahorse Wave 2.4 software.

### Substrate consumption

To evaluate glucose and glutamine uptake, 0.5×106 cells were seeded in 6 wells plate. Media were collected 48h later and immediately frozen at −80°C until analysis. Glutamine concentration was determined as described below with the addition of standard samples for calibration (ranging from 0.01 to 1000 µM). Glucose was measured in cell medium using the Roche diagnostic kit on a Cobas 8000 (Roche Diagnostics) as described previously. Glutamine and glucose consumption was calculated after substraction of the substrate concentration in control medium minus medium with cells for 48h.

### Metabolomic and fluxomic experiments

To investigate intracellular fluxes, 1×10^6^ cells were plated in medium supplemented with 17.5 mM glucose (unlabeled glucose or ^13^C_6_-glucose [CLM-1396, Cambridge Isotope Laboratories]) or 2 mM glutamine (unlabeled glutamine or ^13^C_5_-glutamine [CLM-1822]) and incubated for 5h at 37 °C. Cell pellets were immediately frozen at −80 °C until analysis. Proteins were analyzed for ^13^C labeling of organic acid metabolites and ^13^C labeling of amino acid fraction by LC-HRMS. Data acquisition and processing were achieved using MassLynx and TargetLynx softwares, respectively. Both labeled (from m+1 to m+6) and unlabeled (m+0) compounds were detected by their calculated theoretical *m/z* with a mass tolerance of ± 0.001 Da. The amount of each isotopologue was used to calculate the total amount of each metabolite. ^13^C-enrichments were assessed using IsoCor software (Millard *et al*., 2019).

### Orthotopic GBM mice model

Immunodeficient NSG (NOD.Cg-PrkdcscidIl2rgtm1Wjl/SzJ) mice were housed in the animal facility of the University of Nantes (UTE, SFR F. Bonamy) with support from the LabEx IGO and The Jackson Laboratory (Charles River Lab). Animals were housed under specific pathogen-free status and used between 8 to 12 weeks of age, accordingly to institutional guidelines (Agreement N°17166; Regional ethics committee of the Pays de la Loire; France). GBM cells (1×104 in 2µL PBS) were orthotopically injected in the subventricular zone as previously described (Chauvin *et al*., 2019). Animals were observed daily and euthanized when clinical signs of pain occurring. Brains were collected and fixed in cold PFA 4%. Brain sections (2µm) were used for immunohistochemistry using anti-human MHC class I antibody (clone EPR1394Y; Abgent, CA). Slides were scanned using Nanozoomer (Hamamatsu) and images were analyzed using NPD view 2.3 software.

In order to isolate *in vivo* radiation-escaping cells, mice brain hemisphere bearing the tumor was irradiated 7 days after tumor cell injection at 15Gy using the XRAD225Cx small animal irradiator (Precision XRay Inc.). Tumor cells were isolated using Adult Brain Dissociation kit (130-107-677, Milteniy Biotec) from mice brain at day 20 and day 35 and cultured in NSC medium.

### Statistical analysis

Results are presented as mean ± SD from at least three independent experiments. For statistical analyses, Student t-test, one-way or two-way ANOVA were performed using Prism 7.0 GraphPad Software. P-value below 0.05 was considered significant. On graph comparing control group to CRISPR-Cas9 groups, the p-value for the least significant group is indicated.

## Acknowledgements

This work was supported by Ministère de la Recherche, la Ligue contre le cancer (Comité de Loire Atlantique and Comité de Vendée) and Fondation ARC. We thank core facilities GenoBIRD, Micropicell, Cytocell, Mass Spectrometry of SFR Bonamy and the biochemistry laboratory of Nantes CHU. We thank the LabEX IGO program supported by the National Research Agency (ANR-11-LABX-0016-01) for providing animals. We also thank Audrey Aguesse from the Mass Spectrometry Core Facility of Nantes University for expert technical assistance.

## Notes

**Competing Interest Statement:** The authors declare no conflict of interest

### Competing Interest Statement

The authors have declared no competing interest.

### Summary of Updates

this is not a revised version

## References

Alzial, G. et al. (2022) ‘Wild-type isocitrate dehydrogenase under the spotlight in glioblastoma’, Oncogene, 41(5), pp. 613–621. Available at: https://doi.org/10.1038/s41388-021-02056-1.

Bao, S. et al. (2006) ‘Glioma stem cells promote radioresistance by preferential activation of the DNA damage response’, Nature, 444(7120), pp. 756–760. Available at: https://doi.org/10.1038/nature05236.

Cappel, D.A. et al. (2019) ‘Pyruvate-Carboxylase-Mediated Anaplerosis Promotes Antioxidant Capacity by Sustaining TCA Cycle and Redox Metabolism in Liver’, Cell Metabolism, 29(6), pp. 1291-1305.e8. Available at: https://doi.org/10.1016/j.cmet.2019.03.014.

Chauvin, C. et al. (2019) ‘NKG2D Controls Natural Reactivity of Vγ9Vd2 T Lymphocytes against Mesenchymal Glioblastoma Cells’, Clinical Cancer Research: An Official Journal of the American Association for Cancer Research, 25(23), pp. 7218–7228. Available at: https://doi.org/10.1158/1078-0432.CCR-19-0375.

Chen, J. et al. (2012) ‘A restricted cell population propagates glioblastoma growth after chemotherapy’, Nature, 488(7412), pp. 522–526. Available at: https://doi.org/10.1038/nature11287.

Chen, R. et al. (2010) ‘A hierarchy of self-renewing tumor-initiating cell types in glioblastoma’, Cancer Cell, 17(4), pp. 362–375. Available at: https://doi.org/10.1016/j.ccr.2009.12.049.

Cheng, T. et al. (2011) ‘Pyruvate carboxylase is required for glutamine-independent growth of tumor cells’, Proceedings of the National Academy of Sciences of the United States of America, 108(21), pp. 8674–8679. Available at: https://doi.org/10.1073/pnas.1016627108.

Davidson, S.M. et al. (2016) ‘Environment Impacts the Metabolic Dependencies of Ras-Driven Non-Small Cell Lung Cancer’, Cell Metabolism, 23(3), pp. 517–528. Available at: https://doi.org/10.1016/j.cmet.2016.01.007.

Dirkse, A. et al. (2019) ‘Stem cell-associated heterogeneity in Glioblastoma results from intrinsic tumor plasticity shaped by the microenvironment’, Nature Communications, 10(1), p. 1787. Available at: https://doi.org/10.1038/s41467-019-09853-z.

Fan, T.W.M. et al. (2009) ‘Altered regulation of metabolic pathways in human lung cancer discerned by (13)C stable isotope-resolved metabolomics (SIRM)’, Molecular Cancer, 8, p. 41. Available at: https://doi.org/10.1186/1476-4598-8-41.

Garnier, D. et al. (2019) ‘Glioblastoma Stem-Like Cells, Metabolic Strategy to Kill a Challenging Target’, Frontiers in Oncology, 9, p. 118. Available at: https://doi.org/10.3389/fonc.2019.00118.

Gaude, E. et al. (2018) ‘NADH Shuttling Couples Cytosolic Reductive Carboxylation of Glutamine with Glycolysis in Cells with Mitochondrial Dysfunction’, Molecular Cell, 69(4), pp. 581-593.e7. Available at: https://doi.org/10.1016/j.molcel.2018.01.034.

Hanahan, D. and Weinberg, R.A. (2011) ‘Hallmarks of cancer: The next generation’, Cell, 144(5), pp. 646–674. Available at: https://doi.org/10.1016/j.cell.2011.02.013.

Jin, X. et al. (2017) ‘Targeting glioma stem cells through combined BMI1 and EZH2 inhibition’, Nature Medicine, 23(11), pp. 1352–1361. Available at: https://doi.org/10.1038/nm.4415.

Lao-On, U. et al. (2020) ‘c-Myc directly targets an over-expression of pyruvate carboxylase in highly invasive breast cancer’, Biochimica et Biophysica Acta (BBA) - Molecular Basis of Disease, 1866(3), p. 165656. Available at: https://doi.org/10.1016/j.bbadis.2019.165656.

Lathia, J.D. et al. (2015) ‘Cancer stem cells in glioblastoma’, Genes & Development, 29(12), pp. 1203–1217. Available at: https://doi.org/10.1101/gad.261982.115.

Lee, J. et al. (2006) ‘Tumor stem cells derived from glioblastomas cultured in bFGF and EGF more closely mirror the phenotype and genotype of primary tumors than do serum-cultured cell lines’, Cancer Cell, 9(5), pp. 391–403. Available at: https://doi.org/10.1016/j.ccr.2006.03.030.

Lin, Q. et al. (2020) ‘Targeting Pyruvate Carboxylase by a Small Molecule Suppresses Breast Cancer Progression’, Advanced Science, 7(9). Available at: https://doi.org/10.1002/advs.201903483.

Lottaz, C. et al. (2010) ‘Transcriptional profiles of CD133+ and CD133-glioblastoma-derived cancer stem cell lines suggest different cells of origin’, Cancer Research, 70(5), pp. 2030–2040. Available at: https://doi.org/10.1158/0008-5472.CAN-09-1707.

Ma, M.-Z. et al. (2016) ‘Long Noncoding RNA GCASPC, a Target of miR-17-3p, Negatively Regulates Pyruvate Carboxylase-Dependent Cell Proliferation in Gallbladder Cancer’, Cancer Research, 76(18), pp. 5361–5371. Available at: https://doi.org/10.1158/0008-5472.CAN-15-3047.

Metallo, C.M. et al. (2011) ‘Reductive glutamine metabolism by IDH1 mediates lipogenesis under hypoxia’, Nature [Preprint]. Available at: https://doi.org/10.1038/nature10602.

Meyer, M. et al. (2015) ‘Single cell-derived clonal analysis of human glioblastoma links functional and genomic heterogeneity’, Proceedings of the National Academy of Sciences of the United States of America, 112(3), pp. 851–856. Available at: https://doi.org/10.1073/pnas.1320611111.

Millard, P. et al. (2019) ‘IsoCor: isotope correction for high-resolution MS labeling experiments’, Bioinformatics, 35(21), pp. 4484–4487. Available at: https://doi.org/10.1093/bioinformatics/btz209.

Mullen, A.R. et al. (2012) ‘Reductive carboxylation supports growth in tumour cells with defective mitochondria’, Nature, 481(7381), pp. 385–388. Available at: https://doi.org/10.1038/nature10642.

Oizel, K. et al. (2017) ‘Efficient Mitochondrial Glutamine Targeting Prevails Over Glioblastoma Metabolic Plasticity’, Clinical Cancer Research: An Official Journal of the American Association for Cancer Research, 23(20), pp. 6292–6304. Available at: https://doi.org/10.1158/1078-0432.CCR-16-3102.

Patel, A.P. et al. (2014) ‘Single-cell RNA-seq highlights intratumoral heterogeneity in primary glioblastoma’, Science (New York, N.Y.), 344(6190), pp. 1396–1401. Available at: https://doi.org/10.1126/science.1254257.

Phannasil, P. et al. (2015) ‘Pyruvate Carboxylase Is Up-Regulated in Breast Cancer and Essential to Support Growth and Invasion of MDA-MB-231 Cells’, PloS One, 10(6), p. e0129848. Available at: https://doi.org/10.1371/journal.pone.0129848.

Sellers, K. et al. (2015) ‘Pyruvate carboxylase is critical for non–small-cell lung cancer proliferation’, The Journal of Clinical Investigation, 125(2), pp. 687–698. Available at: https://doi.org/10.1172/JCI72873.

Singh, S.K. et al. (2004) ‘Identification of human brain tumour initiating cells’, Nature, 432(7015), pp. 396–401. Available at: https://doi.org/10.1038/nature03128.

Sottoriva, A. et al. (2013) ‘Intratumor heterogeneity in human glioblastoma reflects cancer evolutionary dynamics’, Proceedings of the National Academy of Sciences of the United States of America, 110(10), pp. 4009–4014. Available at: https://doi.org/10.1073/pnas.1219747110.

Stupp, R. et al. (2009) ‘Effects of radiotherapy with concomitant and adjuvant temozolomide versus radiotherapy alone on survival in glioblastoma in a randomised phase III study: 5-year analysis of the EORTC-NCIC trial’, The Lancet Oncology, 10(5), pp. 459–466. Available at: https://doi.org/10.1016/S1470-2045(09)70025-7.

Tang, X. et al. (2021) ‘Targeting Glioblastoma Stem Cells: A Review on Biomarkers, Signal Pathways and Targeted Therapy’, Frontiers in Oncology, 11. Available at: https://www.frontiersin.org/article/10.3389/fonc.2021.701291 (Accessed: 21 April 2022).

Tardito, S. et al. (2015) ‘Glutamine synthetase activity fuels nucleotide biosynthesis and supports growth of glutamine-restricted glioblastoma’, Nature Cell Biology, 17(12), pp. 1556–1568. Available at: https://doi.org/10.1038/ncb3272.

Verhaak, R.G.W. et al. (2010) ‘Integrated Genomic Analysis Identifies Clinically Relevant Subtypes of Glioblastoma Characterized by Abnormalities in PDGFRA, IDH1, EGFR, and NF1’, Cancer Cell, 17(1), pp. 98–110. Available at: https://doi.org/10.1016/j.ccr.2009.12.020.

Wang, J. et al. (2008) ‘CD133 negative glioma cells form tumors in nude rats and give rise to CD133 positive cells’, International Journal of Cancer, 122(4), pp. 761–768. Available at: https://doi.org/10.1002/ijc.23130.

Wang, Q. et al. (2017) ‘Tumor Evolution of Glioma-Intrinsic Gene Expression Subtypes Associates with Immunological Changes in the Microenvironment’, Cancer Cell, 32(1), pp. 42-56.e6. Available at: https://doi.org/10.1016/j.ccell.2017.06.003.

Wise, D.R. et al. (2011) ‘Hypoxia promotes isocitrate dehydrogenase-dependent carboxylation of α-ketoglutarate to citrate to support cell growth and viability’, Proceedings of the National Academy of Sciences of the United States of America, 108(49), pp. 19611–19616. Available at: https://doi.org/10.1073/pnas.1117773108.

