## Supplementary Figures for "Pyruvate carboxylation identifies Glioblastoma Stem-like Cells opening new metabolic strategy to prevent tumor recurrence"

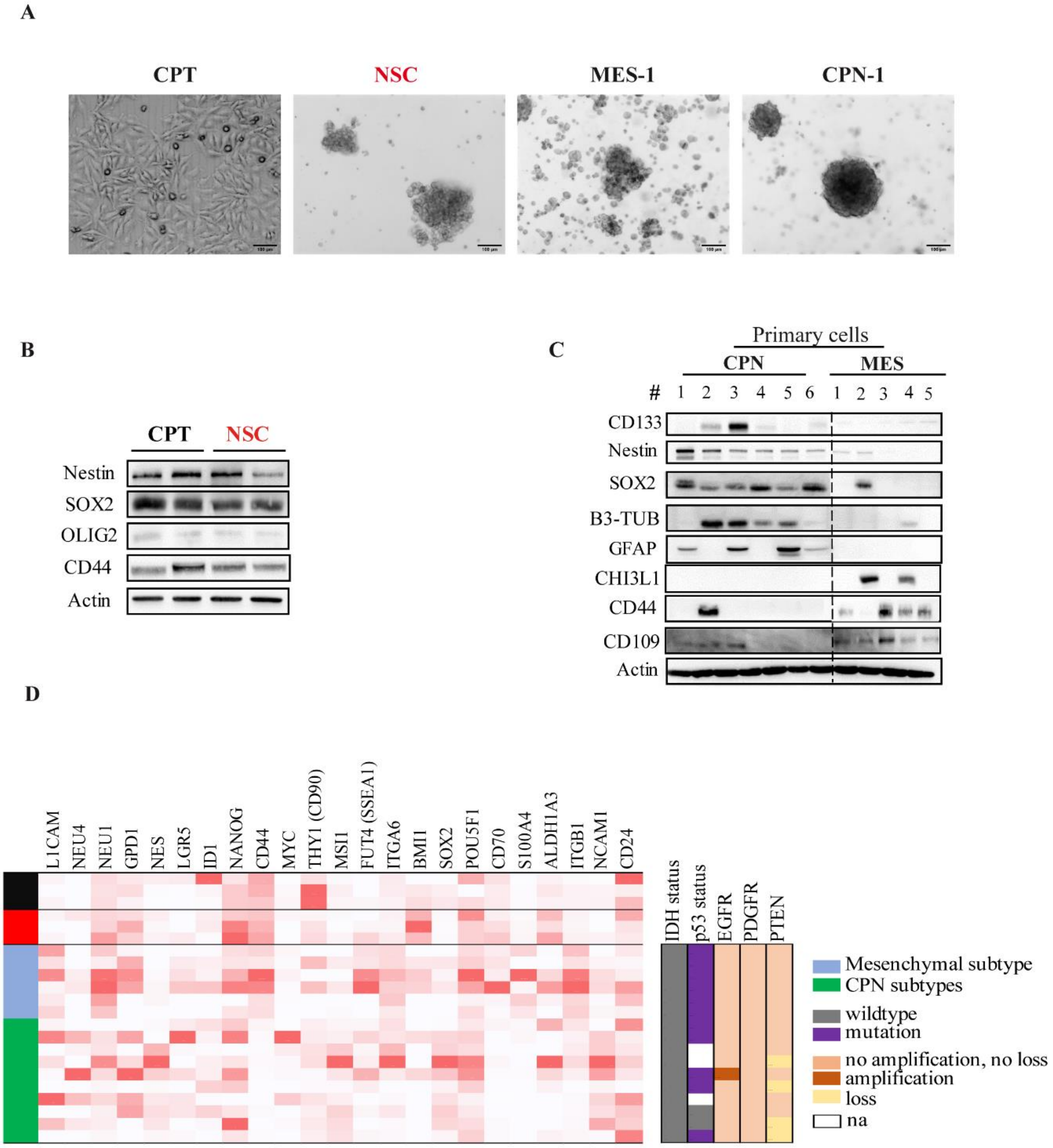


**Supplementary Figure 1:**

**A**. Representative pictures of U251 cells cultured in complete (CPT) or NSC media, as well as patient-derived primary cultures in NSC medium depending on their molecular subtype. **B-C**. Representative immunoblots of stemness, differentiation and aggressiveness markers expression in U251 cells according to culture conditions (B) and in patient-derived primary cultures depending on their molecular subtype (C). **D**. Heatmap representing transcriptomic expression of diverse stemness markers (*left panel*) in U251 cells depending on the culture conditions, as well as in patient-derived primary cultures depending on their molecular subtypes. U251 cells cultured in CPT media are indicated in black, in CPT in red whereas MES primary GBM cells are indicated in blue and CPN in green. Main genetic alterations are also indicated for patient-derived primary cultures (*right panel*) such as IDH and p53 mutation status, and amplification or loss of EGFR, PDGFR and PTEN.

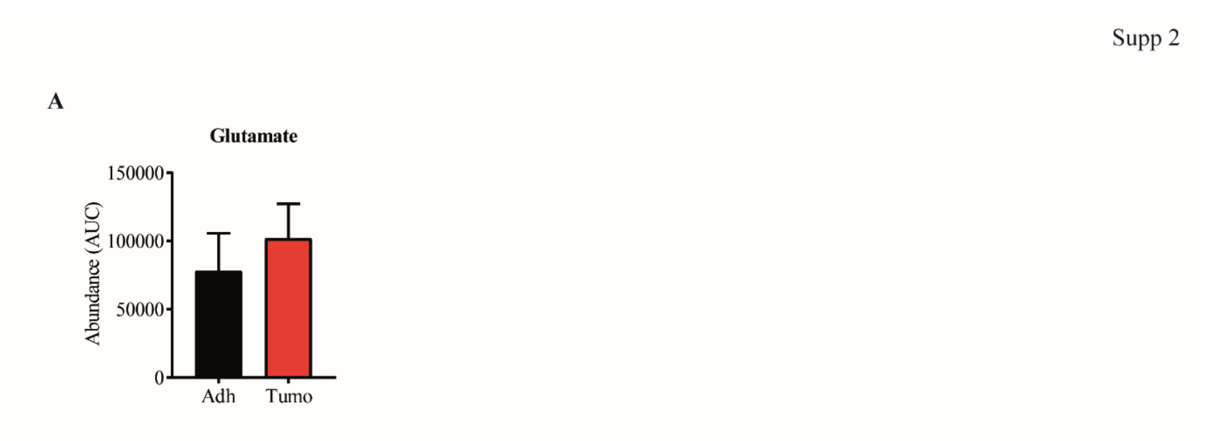


**Supplementary Figure 2:**

**A**. Intracellular abundance of glutamate in U251 cells depending on culture conditions by mass spectrometry. Student t-test (n=3).

Supp 3

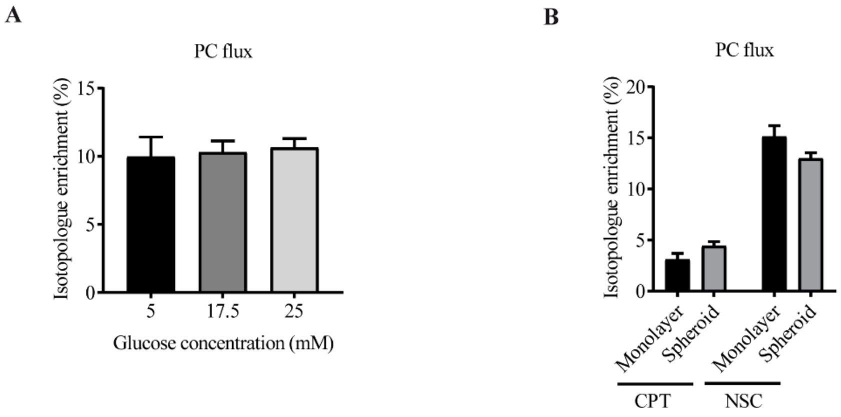


**Supplementary Figure 3:**

**A**. PC activity is not affected by glucose concentration in the media in U251 cells. PC activity was determined by the enrichment of malate m+3 isotopologue in U251 cells cultured in NSC medium. **B**. Cell culture as spheroids does not affect PC activity in all cells, independently of the original growth medium. PC activity was determined in U251 cells cultured in complete medium (CPT) or Neural stem cells medium (NSC) PC activity by the enrichment of malate m+3 isotopologue.

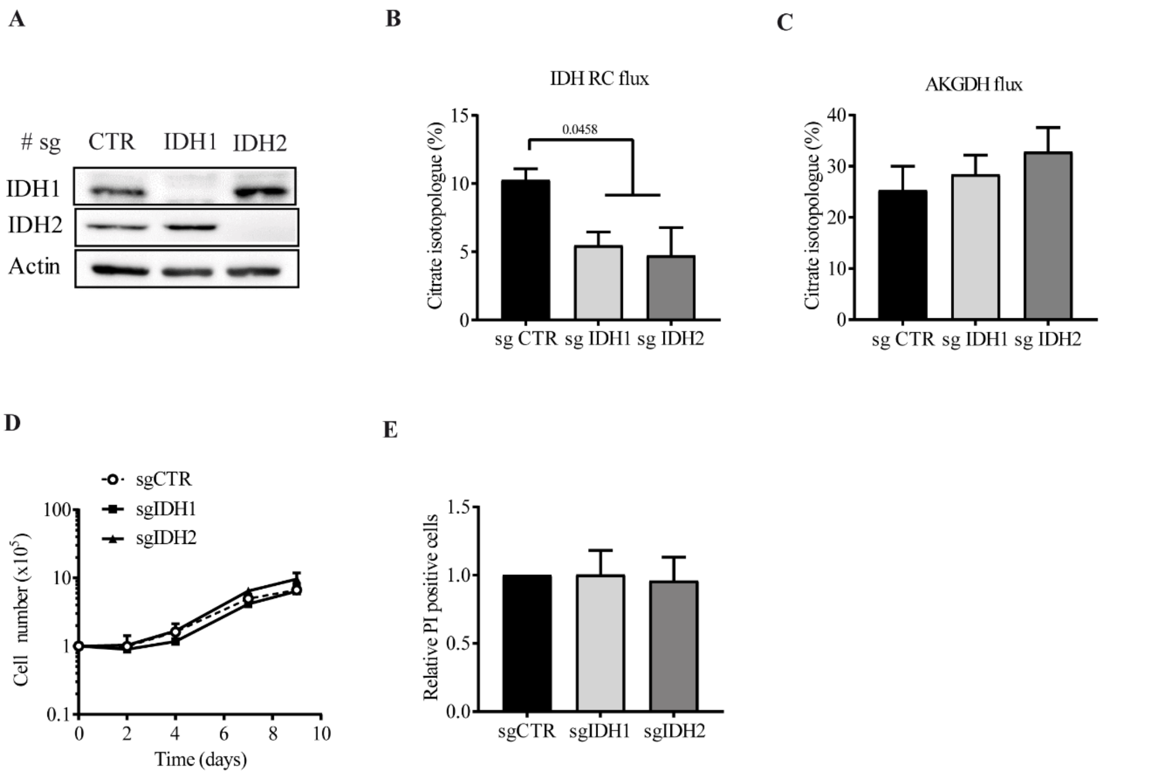


**Supplementary Figure 4:**

**A**. Representative immunoblot of IDH1 and IDH2 silencing in tumoroid U251 cells. IDH 1/2 were silenced in tumoroid U251 cells. **B-C**. IDH (B) and AKGDH (C) activities assessed by (m+5) and (m+4) citrate enrichment respectively, following culture with $^{13}C_5$-labeled glutamine. (n=4). **D**. Cell proliferation of tumoroid cells following IDH silencing. Cell proliferation was assessed over 9 days by manual cell counting following eosin staining (n=3). **E**. Relative cell death measured by flow cytometry following PI staining after IDH silencing. One way ANOVA (n=3).

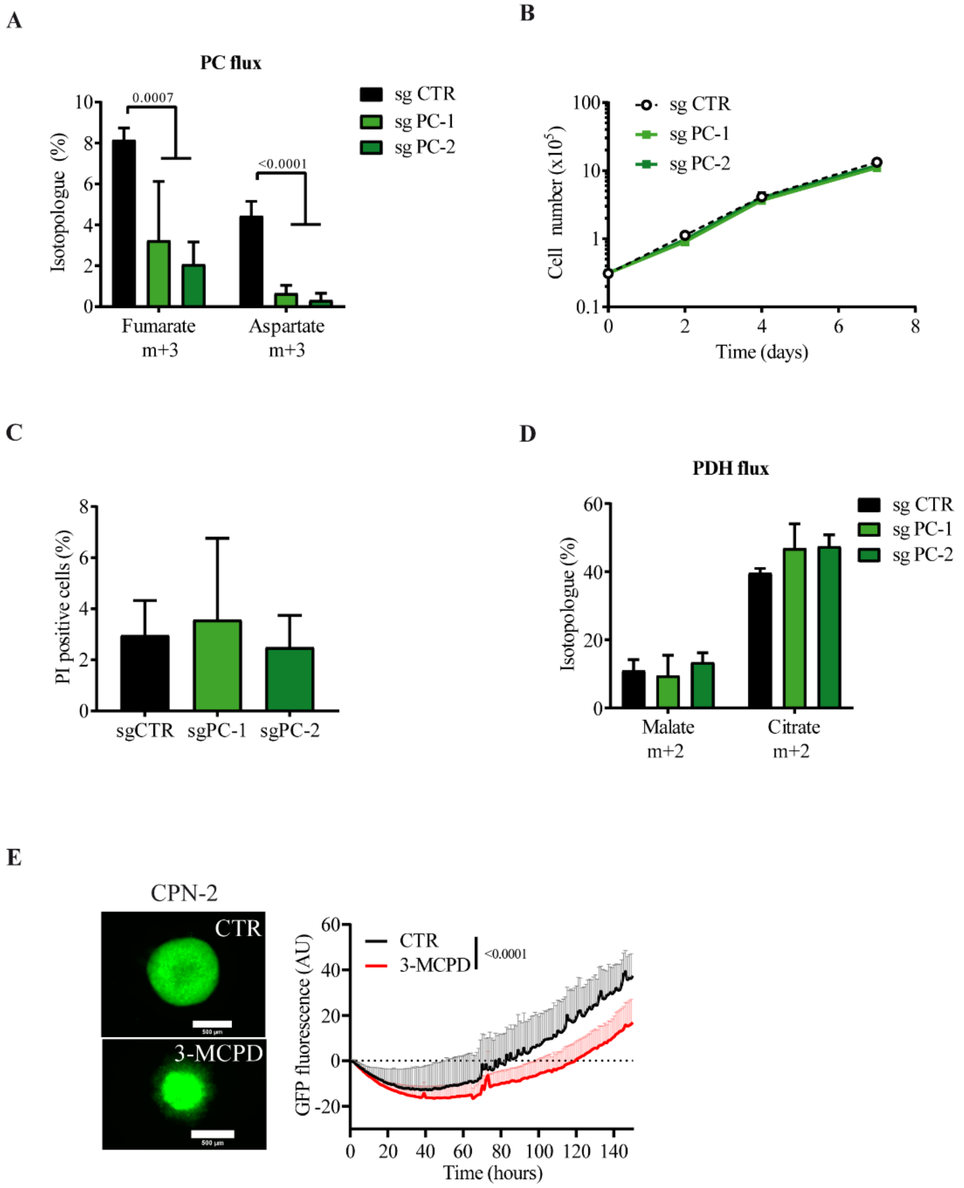


**Supplementary Figure 5:**

**A**. Glycolytic-derived (m+3) isotopologue enrichment in PC-silenced cells. Enrichment of (m+3) fumarate and aspartate were determined by mass spectrometry following culture with $^{13}C_6$-labeled glucose. Two-way ANOVA (n=4). **B**. Proliferation of control and PC silenced in adherent U251 cells over 9 days. Student t-test (n=3). **C**. Cell death measured by flow cytometry following PI staining in adherent U251 cells according to PC status. Two-way ANOVA (n=3). **D**. Glycolytic-derived (m+2) isotopologue enrichment in PC-silenced cells. Enrichment of (m+2) malate and citrate were determined by mass spectrometry following culture with $^{13}C_6$-labeled glucose. Two-way ANOVA (n=4). **E**. Proliferation of GFP-CPN-2 cells following 3-MCPD treatment. Fluorescence intensity was followed over 8 days with Incucyte device (Objective 4X, scale bar = 500µm). Two-way ANOVA (n=3).

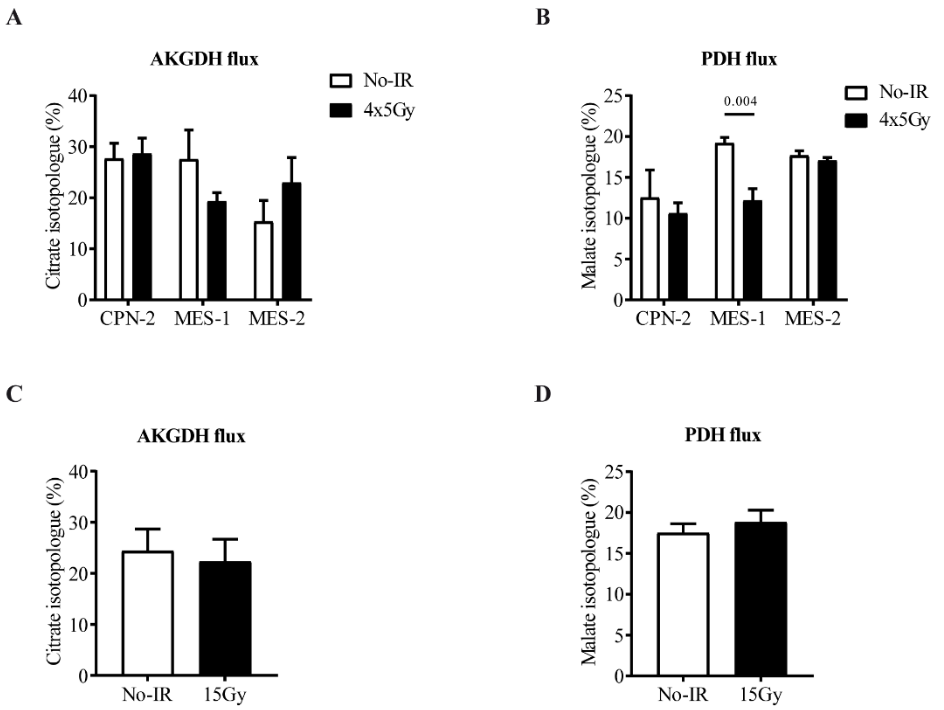


**Supplementary Figure 6:**

**A.** AKGDH activity in 4x5Gy radiation-escaping cells determined by (m+4) citrate enrichment following culture with $^{13}C_5$-labeled glutamine. AKGDH activity was assessed in the 3 primary cultures giving rise to radiation-surviving cells. Student t-test (n=3). **B**. PDH activity in 4x5Gy radiation-escaping cells determined by (m+2) $^{13}C$-malate enrichment by mass spectrometry following culture with $^{13}C_6$-labeled glucose. PDH activity was assessed in the 3 primary cultures giving rise to radiation-surviving cells. Student t-test (n=3). **C.** AKGDH activity in 15Gy radiation-escaping cells determined by (m+4) citrate enrichment following culture with $^{13}C_5$-labeled glutamine. **D**. PDH activity in 15Gy radiation-escaping cells determined by (m+2) $^{13}C$-malate enrichment by mass spectrometry following culture with $^{13}C_6$-labeled glucose.
